# Regulation of IFNγ production by ZFP36L2 in T cells is context-dependent

**DOI:** 10.1101/2024.02.09.579641

**Authors:** Nordin D. Zandhuis, Aurélie Guislain, Abeera Popalzij, Sander Engels, Branka Popović, Martin Turner, Monika C. Wolkers

## Abstract

CD8^+^ T cells kill target cells by releasing cytotoxic molecules and pro-inflammatory cytokines, such as TNF and IFNγ. The magnitude and duration of cytokine production is defined by post-transcriptional regulation, and a critical regulator herein are RNA-binding proteins (RBPs). Although the functional importance of RBPs in regulating cytokine production is established, the kinetics and mode of action through which RBPs control cytokine production is not well understood. Previously, we showed that the RBP ZFP36L2 blocks translation of pre-formed cytokine encoding mRNA in quiescent memory T cells. Here, we uncover that ZFP36L2 regulates cytokine production in a context-dependent manner. T cell-specific deletion of ZFP36L2 (CD4-cre) had no effect on T cell development, or on cytokine production during early time points (2-6h) of T cell activation. In contrast, ZFP36L2 specifically dampened the production of IFNγ during prolonged T cell activation (20-48h). ZFP36L2 deficiency also resulted in increased production of IFNγ production in tumour-infiltrating T cells that are chronically exposed to antigen. Mechanistically, ZFP36L2 regulates IFNγ production at late time points of activation by destabilizing *Ifng* mRNA in an AU-rich element-dependent manner. Together, our results reveal that ZFP36L2 employs different regulatory nodules in effector and memory T cells to regulate cytokine production.

## INTRODUCTION

T cells are essential in protecting the host from pathogenic insults and malignancies. When exposed to their cognate antigen by antigen-presenting cells (APCs), naïve T cells are primed in secondary lymphoid organs, they differentiate and produce effector molecules to kill target cells. Upon target cell clearance, the effector T cell population contracts, and a small percentage of antigen-specific T cells persists as memory T cells (1, 2).

T cell differentiation into effector cells requires the reorganization of gene expression instructed by dynamic epigenetic, transcriptional, post-transcriptional and metabolic changes (3–5). Many factors influence these alterations in gene expression, such as TCR affinity to the antigen, antigen load, costimulatory molecule expression, and the presence of metabolites, chemokines, and cytokines (6, 7). The effector function of T cells also requires tight gene regulation, including cytokine production (8–10). Excessive cytokine production correlates with the development of autoimmune disorders (11–13) and organ dysfunction in COVID-19 patients (14). Conversely, insufficient production of the pro-inflammatory cytokines Tumor Necrosis Factor (TNF) and Interferon gamma (IFNγ) impedes the clearance of virally infected and tumour cells (15–18). Therefore, to produce adequate levels of effector molecules, the fine-tuning of cytokine production is key. Post-transcriptional regulation by RNA-binding proteins (RBPs) is a critical factor in regulating the intricate control of cytokine expression (19–21).

Over 2000 RBPs are expressed in mammalian cells, which instruct the fate of the mRNA and thus the cellular fate (22). In T cells, the ZFP36 protein family, consisting of ZFP36, ZFP36L1 and ZFP36L2, regulates T cell development, differentiation and effector function (23–27). ZFP36 proteins interact with AU-rich elements (AREs) in the 3’Untranslated regions (3’UTRs) of their target mRNAs (28, 29). The combined loss of ZFP36L1 and ZFP36L2 perturbs thymic T cell development (30), and double deletion of ZFP36 and ZFP36L1 drives CD8^+^ T cell differentiation towards effector cells (23, 25). Also, cytokine production is modulated by ZFP36 proteins (26). ZFP36-deficient murine CD4^+^ T cells produce more IFNγ,TNF, IL-17, and IL-2 (23, 25, 31–33). Likewise, human effector T cells devoid of ZFP36L1 show increased and prolonged production of IL-2, TNF, and IFNγ (34). Conversely, ZFP36L2 prevents the premature translation of pre-formed cytokine mRNA in quiescent memory T cells (35). Whether ZFP36L2 also regulates the production of cytokines during other T cell states, e.g. upon T cell priming or T cell activation is to date not known.

In this study, we investigated the role of ZFP36L2 during T cell differentiation and T cell activation. Whereas T cell development and T cell priming were not affected by ZFP36L2 deficiency, we found that ZFP36L2 represses cytokine production in particular at later time points of T cell activation, with the largest effects on IFNγ. ZFP36L2 deficiency also augments the IFNγ production in tumour-infiltrating T cells (TILs). Intriguingly, and contrary to its role in suppressing translation in memory T cells, ZFP36L2 induces *Ifng* mRNA degradation in an AU-rich element-dependent manner in TILs. PD-1 blockade acted synergistically with ZFP36L2 in driving the expression of IFNγ during continuous T cell activation. We here thus uncover a context-dependent mode of action of ZFP36L2 in regulating T cell function.

## RESULTS

### Normal development and differentiation of ZFP36L2-deficient T cells

ZFP36L2 deficiency has been primarily investigated in combination with other ZFP36 family members (23, 24, 30, 36). To specifically decipher the role of ZFP36L2 in T cells, we used conditional CD4-Cre ZFP36L2 knock-out mice (ZFP36L2^CKO^). Compared to wild type (WT) *Zfp36l2^fl/f^ CD4-Cre^-/-^* littermates, the weight and appearance (posture and agility) of ZFP36L2^CKO^ mice was normal until the end point of measurements, i.e. up to 9-12 months of age (**Supp. Fig. 1A**). The distribution of thymic precursor T cell populations as defined by CD4/CD8 expression and CD44/CD25 expression on double negative (DN) cells was equal between ZFP36L2^CKO^ and WT mice (**Fig. 1A**). This was also true for thymic CD25^-^ Foxp3^+^ Treg precursors and mature CD25^+^ Foxp3^+^ Tregs (**Fig. 1B**). Thus, at later stages of T cell development when ZFP36L2 deficiency is established, T cell development is not affected.

**Figure 1.**
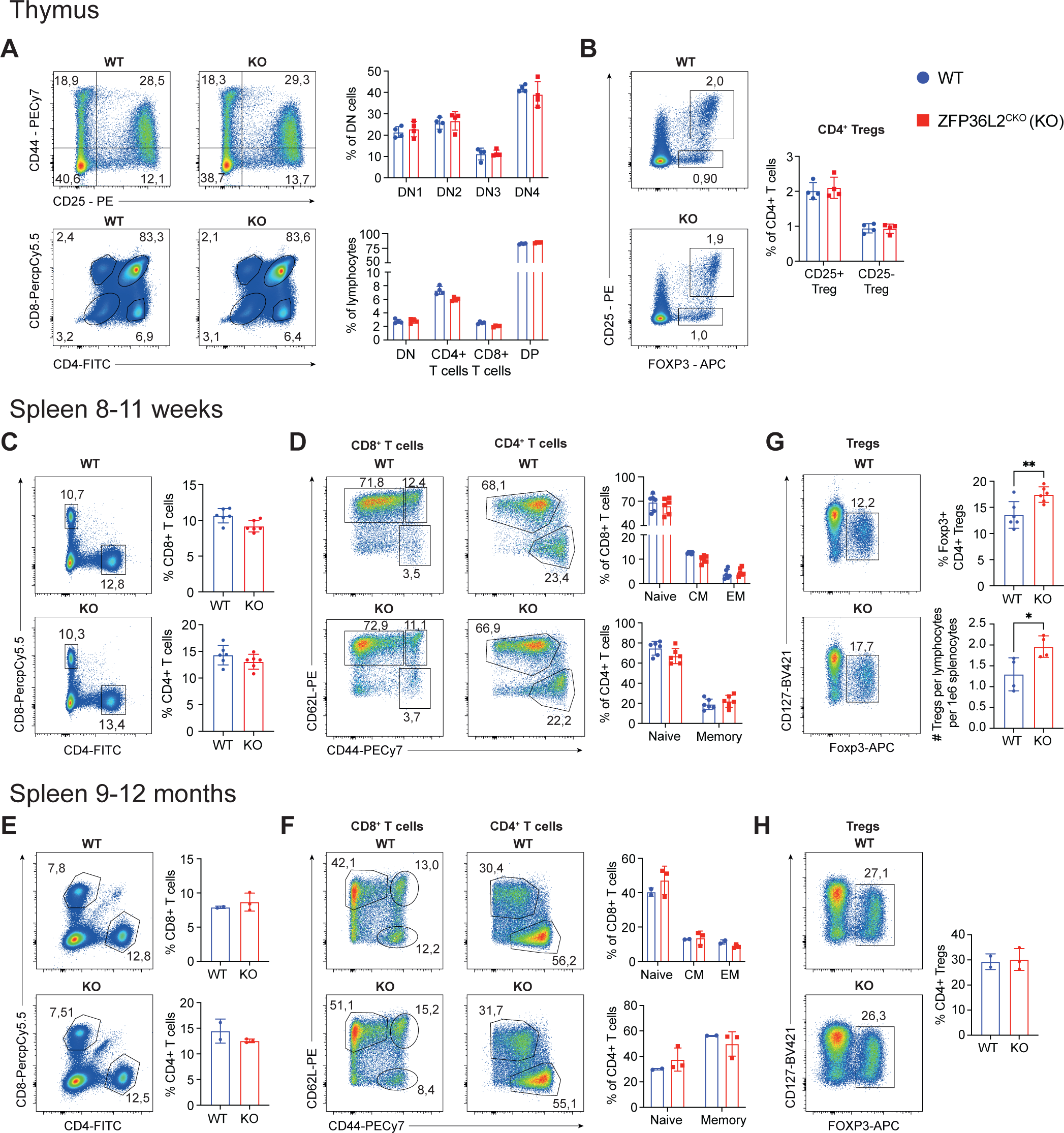
Phenotyping WT and ZFP36L2^CKO^ mice under steady state conditions. **(A-B)** Flow cytometry analysis of bulk thymocytes (A) and thymic Tregs (B) of WT (CD4-cre^-^ *Zfp36l2^fl/fl^*) and ZFP36L2^CKO^ (KO) (CD4-cre^+^ *Zfp36l2^fl/fl^*) mice. n=4/group. **(C-H)** Representative flow cytometry plots and frequencies of splenic WT and KO CD8^+^ and CD4^+^ T cells (C,F), and their respective differentiation status (D,E,G,H), including; naïve (CD62L^+^ CD44^-^), central memory (CD62L^+^ CD44^+^), effector memory (CD62L^-^ CD44^+^) CD8^+^, and naïve (CD62L^+^ CD44^low^), memory (CD62L^-^ CD44^hi^) and regulatory (CD127^lo^ Foxp3^+^) CD4^+^ T cell populations. C-E: n=6 mice/group. F-H: n=2 for WT mice and n=3 for KO mice. Data were analyzed by one-way ANOVA with Tukey multiple comparison correction, mean ± s.d. (**A,B,D,G**), or by two-sided, unpaired Student’s *t*-test, mean ± s.d. (**C,E,F,H**); *p<0.05, **p<0.01.

Under steady state conditions, total cell numbers of spleens were identical in young (8-11 weeks old) ZFP36L2^CKO^ and WT mice and in spleens and livers in middle-aged (9-12 months old) mice, with a slight increase in inguinal lymph nodes (iLNs) (**Supp. Fig. 1B**). The percentage and differentiation status (naïve, effector, memory) of CD8^+^ T cells and CD4^+^ T cells was also unaltered in spleen and iLN (**Fig. 1C-F**, **Supp. Fig. 1C-D**). Frequencies of liver CD4^+^ and CD8^+^ T cells were slightly lower in middle-aged ZFP36L2^CKO^ mice, with no or only limited changes in the T cell differentiation profiles (**Supp. Fig. 1F, G**). Interestingly, the percentage and number of splenic Foxp3^+^ CD127^lo^ regulatory CD4^+^ T cells (Tregs) was ∼1.3 fold increased in young ZFP36L2^CKO^ mice (**Fig. 1G**), which we found originated from thymic and from peripheral Tregs, because the frequency of NRP1-expressing thymic-derived Tregs remained unchanged between splenic WT and ZFP36L2^CKO^ Tregs (**Supp. Fig. 1H**) (37). Treg numbers were also increased in iLNs of middle-aged mice, but this was not evident in the spleen of these mice (**Fig. 1H**, **Supp. Fig. 1G**). Furthermore, FACS-sorted naïve ZFP36L2^CKO^ and WT CD4^+^ T cells differentiated equally well *in vitro* into Tregs upon activation with α-CD3/α-CD28 and recombinant mTGF-β (**Supp. Fig. 1I**), indicating that ZFP36L2 deficiency did not alter the propensity of naïve T cells to differentiate into Tregs. Overall, we did not observe any overt effects during development and at steady state in T cells of ZFP36L2^CKO^ mice.

### ZFP36L2 modulates cytokine production during T cell priming

We next questioned whether ZFP36L2 deficiency influences the cytokine production upon activation of naïve T cells. We crossed ZFP36L2^CKO^ mice with OT-I T cell receptor transgenic mice that recognize the SIINFEKL (N4) peptide of Chicken egg Ovalbumin. FACS-sorted naïve CD44^lo^ WT and ZFP36L2^CKO^ OT-I T cells were co-cultured with peptide-loaded bone marrow-derived dendritic cells (BMDCs). After 20 h and 48 h of T cell priming, we measured the production of IFNγ, TNF, and IL2 upon blocking protein secretion for the last 2 h of activation with Brefeldin A.

At 20h post activation, the percentage of TNF-producing T cells was similar, just like the production of TNF per cell, as determined by its geometric-mean fluorescence intensity (gMFI; **Fig. 2A**). In contrast, the production levels of IFNγ and IL-2 were significantly increased in ZFP36L2^CKO^ OTI T cells compared to WT OTI T cells (**Fig. 2A**). At 48h of activation, the effect of ZFP36L2 was strongest on the production of IFNγ, resulting in a doubling of IFNγ-producing T cells at intermediate (1nM) and low peptide (0.01nM) concentrations, as well as increased IFNγ production per cell (**Fig. 2B**). Similar results were obtained when naïve T cells were primed with the low affinity variant SIIQFEKL (Q4) (**Supp. Fig. 2A, B**). Of note, the antigen threshold for cytokine production upon T cell priming was similar between WT and ZFP36L2^CKO^ OTI T cells (**Fig. 2A, B**). Thus, ZFP36L2 does not alter the threshold for T cell responsiveness upon T cell priming, but it limits the magnitude and duration of IFNγ and IL-2 production.

**Figure 2.**
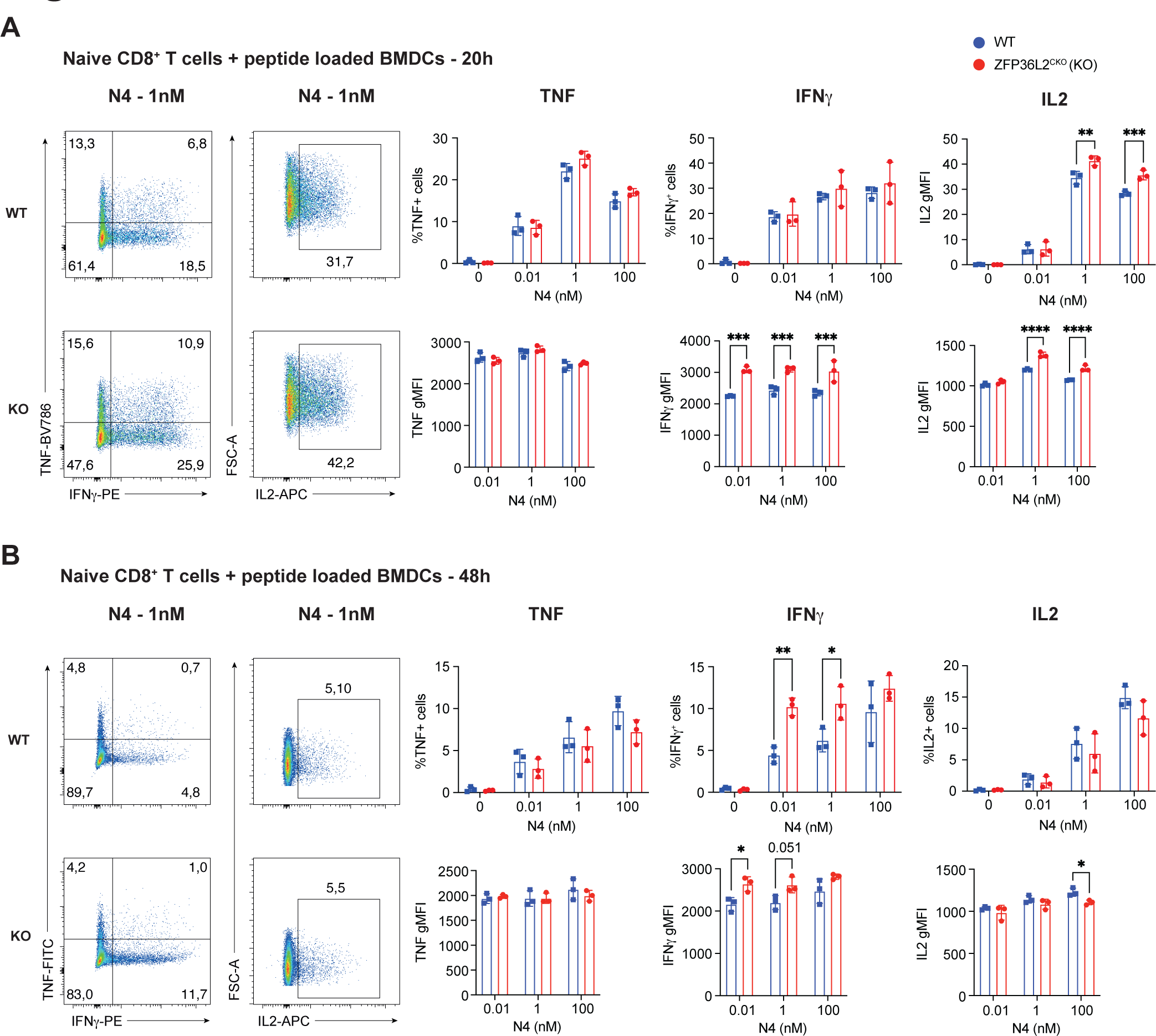
ZFP36L2 regulates IFNγ production during priming of naïve CD8^+^ T cells. **(A,B)** Cytokine production of naïve WT (OT-I^+^ CD4-cre^-^ *Zfp36l2^fl/fl^*) and KO (OT-I^+^ CD4-cre^+^ *Zfp36l2^fl/fl^*) OT-I CD8^+^ T cells that were primed for 20 h (A) or 48 h (B) with bone marrow-derived DCs loaded with indicated concentration of the SIINFEKL (N4) peptide. Brefeldin A was added for the last 2 h of activation. (n=3 mice/group). Percentage (top right) and cytokine production per cell (bottom right) as defined by the Geo-mean fluorescence intensity (gMFI). Data were analyzed by two-way ANOVA with Šidák multiple comparison correction, mean ± s.d. ; *p<0.05; **p<0.01, ***p<0.001, ****p<0.0001.

### Time-specific modulation of IFN**γ** production by ZFP36L2

We then turned to study the role of ZFP36L2 on cytokine production in effector T cells. We activated WT and ZFP36L2^CKO^ OT-I T cells for 24h with MEC.B7.SigOVA cells expressing the N4 peptide and CD80 (38). T cells were then cultured for an additional 3-5 days in the absence of antigen with low IL-7 (hereafter called “rested”) (39). T cell activation for 6h with Ovalbumin-expressing B16-F10 melanoma cells (B16-OVA) only showed a slight, yet significant increase in the per cell production of IFNγ (**Fig. 3A**). Likewise, short term activation (2h and 6h) with different N4 or Q4 peptide concentrations showed a similar antigen threshold for cytokine production, and similar percentages and production levels of all three cytokines for WT and ZFP36L2^CKO^ OT-I T cells (**Supp. Fig. 3A-D**).

**Figure 3.**
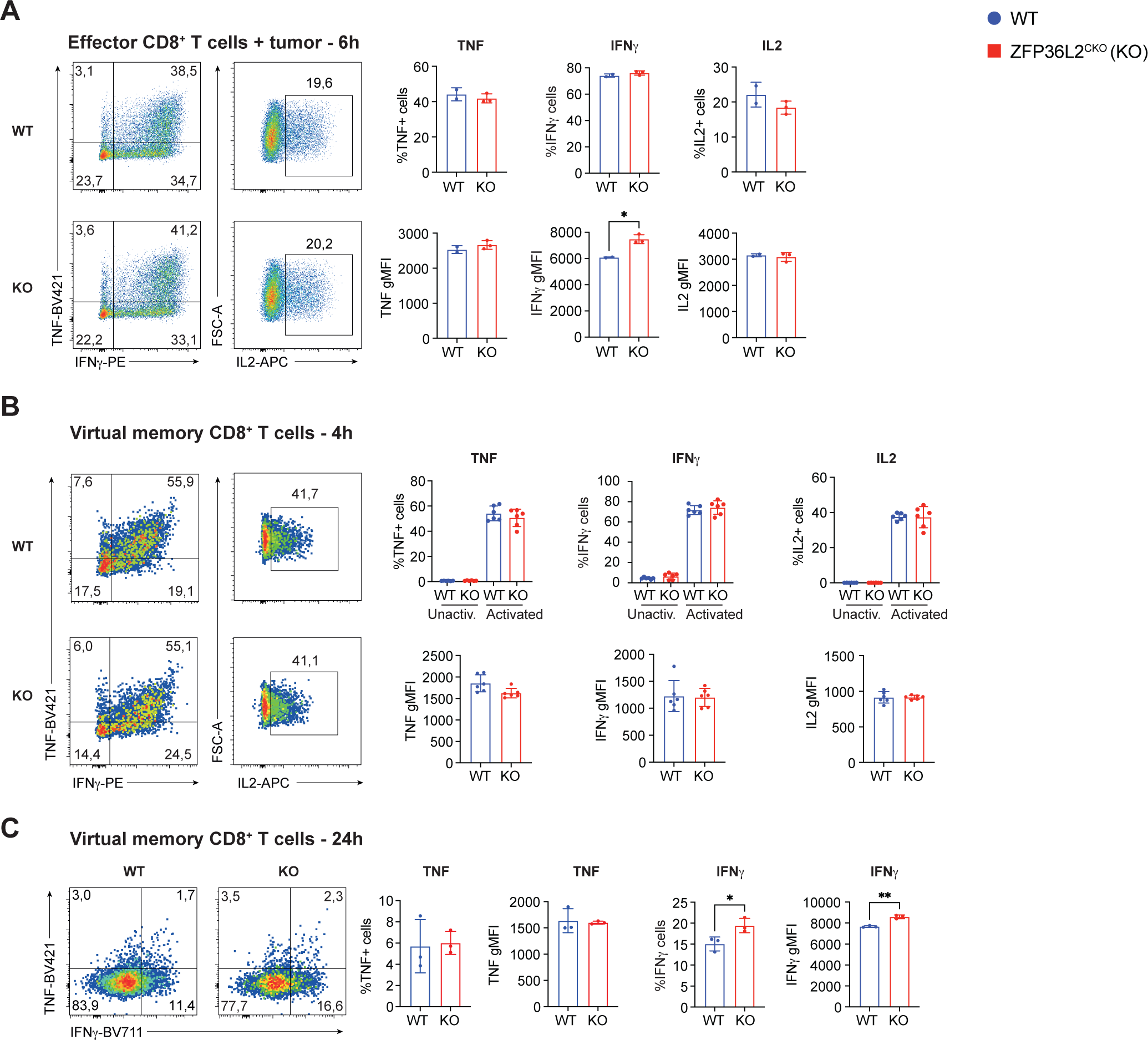
ZFP36L2 dampens IFNγ production in effector CD8^+^ T cells during prolonged activation. **(A)** Cytokine production of rested WT and KO OT-I CD8^+^ T cells after 6 h of co-culture with B16-OVA tumour cells. (n=2 for WT and n=3 KO mice). **(B)** Cytokine production of CD44^hi^ virtual memory (VM) CD8^+^ T cells after 4 h stimulation with PMA/Ionomycin. (n=6 mice/group). **(C)** Cytokine production of CD8^+^ VM T cells after 24 h stimulation with α-CD3 and α-CD28. (n=3 mice/group). In all experiments, Brefeldin A was added for the last 2 h of activation. Graphs display mean and S.D. values. Data were analyzed by two-sided, unpaired Student’s *t*-test, mean ± s.d. ; *p<0.05, **p<0.01.

Like effector T cells, memory T cells rapidly produce cytokines upon reactivation (40). This also holds true for naturally occurring CD44^hi^ virtual memory (VM) T cells (41, 42). In line with our previous observations that ZFP3L2 rapidly releases cytokine mRNA upon reactivation (35), we found no effect of ZFP36L2 deficiency on short-term (4h) reactivation of CD8 VM T cells with phorbol 12-myristate 13-acetate (PMA)/Ionomycin (**Fig. 3B**). In contrast, at 24h of reactivation with αCD3/αCD28, at a time point when the IL-2 production is undetectable and TNF production is low (34, 43), the percentage and magnitude of IFNγ production was significantly increased (1.25 fold) in CD8^+^ ZFP36L2^CKO^ VM cells (**Fig. 3C**). Thus, ZFP36L2 deficiency modulates the production of IFNγ primarily at later time points.

### Zfp36l2 attenuates the frequency of IFN**γ** producing tumour-infiltrating T cells

Because the strongest effect of ZFP36L2 deficiency on IFNγ was observed at later time points of T cell priming and (re)activation (**Fig. 2B**, **Fig. 3C**), we questioned whether ZFP36L2 deficiency also modulates the production of IFNγ when T cells are continuously exposed to antigen. This is for instance the case in the tumour microenvironment, as exemplified in the B16-OVA melanoma model (44).

WT and ZFP36L2^CKO^ effector OT-I T cells were transferred into B16-OVA bearing mice (**Supp. Fig. 4A**). Over the course of 43 days, the tumour growth in this aggressive model was comparable in mice that received WT or ZFP36L2^CKO^ effector OT-I T cells (**Supp. Fig. 4B, C**). Tumour size and recovery of OT-I TILs at day 11 per milligram of tumour B16-OVA tumours was comparable, irrespective of the genetic background of transferred T cells (**Supp. Fig. 4D, E**). Likewise, the expression levels of the co-inhibitory receptor PD-1 were equally high in WT and ZFP36L2^CKO^ TILs (**Supp. Fig. 4F**), as was the frequency of PD-1^+^TIM3^-^ and TCF1^+^ TOX^+^ precursor exhausted TILs **(Supp. Fig. 4F, G**). The differentiation status between quiescent (LY108^+^CD69^+^), cycling (LY108^+^CD69^-^) and terminal dysfunctional (LY108^-^ CD69^+^, PD-1^+^ TIM3^+^) WT and ZFP36L2^CKO^ TILs was also indistinguishable (**Supp. Fig. 4H**), indicating that the differentiation status of TILs was not sensitive to ZFP36L2 deficiency.

Because chronic antigen exposure and the immunosuppressive environment of tumours results in gradual loss of T cell effector function (45, 46), we next questioned if ZFP36L2 deficiency modulated effector molecules in TILs. The expression of the degranulation marker CD107a was identical between WT and ZFP36L2^CKO^ TILs (**Fig. 4B**), and the percentage and production of Granzyme B per cell was slightly, but not significantly elevated in ZFP36L2^CKO^ TILs (**Fig. 4C**). In sharp contrast, even though the IFNγ production per cell was not altered, the percentage of IFNγ producing TILs doubled in ZFP36L2^CKO^ TILs, reaching 36.8±10.4% compared to 16.0±7.1% of WT TILs (**Fig. 4D**). Thus, also in the immunosuppressive tumour environment, ZFP36L2 deficiency results in substantial increases in IFNγ-producing T cells.

**Figure 4.**
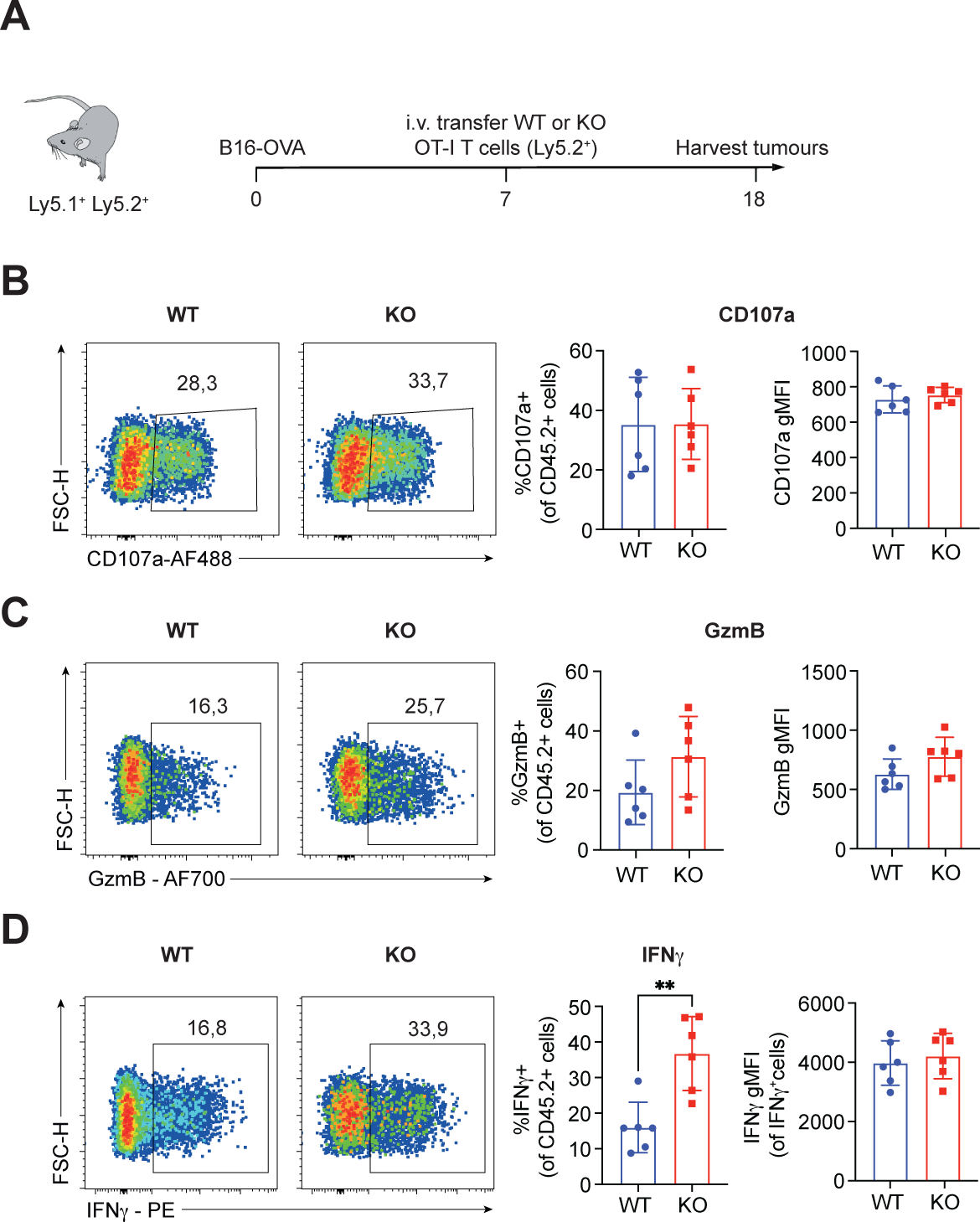
ZFP36L2-deficient tumour-specific T cells display increased production and *Ifng* mRNA stability *ex vivo.* **(A)** Schematic overview of the B16-OVA tumour model. **(B-D)** *Ex vivo* frequency and expression levels of CD107a (B), Granzyme B (C) and IFNγ (D) by WT and KO tumour-specific OT-I T cells after 4 hours of culture in the presence of Brefeldin A and Monensin (n=6 mice/group). Data were analyzed by two-sided, unpaired Student’s *t*-test, mean ± s.d. ; *p<0.05, **p<0.01.

### ZFP36L2 regulates IFN**γ** production through direct interaction with the *Ifng* mRNA

To study the role of ZFP36L2 on IFNγ in more detail, we tested whether our previously established *in vitro* co-culture system of continuous antigen stimulation (47) recapitulated the effect of ZFP36L2 deficiency on IFNγ production. WT and ZFP36L2^CKO^ OT-I effector T cells were exposed to B16-OVA cells for 24h, and then transferred onto freshly seeded B16-OVA cells for another 24h (48h) (**Fig. 5A**). As previously reported (47) and in line with the cytokine production kinetics of T cells (34, 43), the production of TNF and IL-2 dropped to undetectable levels at 24h and 48h of activation. Only the production of IFNγ was sustained (**Fig. 5B**). Importantly, this *in vitro* system mirrored the effect of ZFP36L2 in TILs: the percentage of IFNγ producing cells was substantially higher in ZFP36L2^CKO^ T cells compared to WT T cells (**Fig. 5B**), therefore this model allowed us to decipher how ZFP36L2 modulates the production of IFNγ.

**Figure 5.**
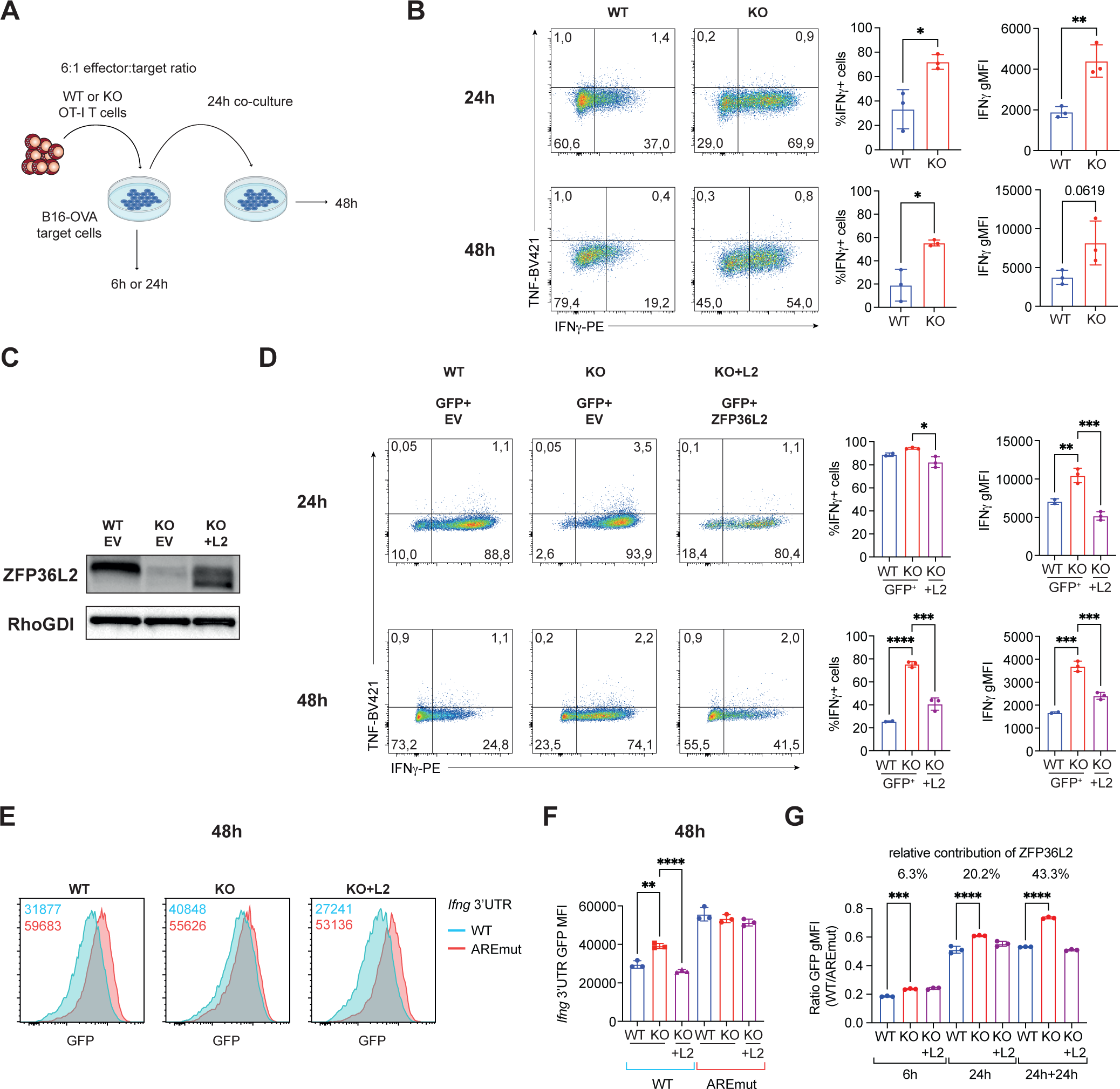
ZFP36L2 regulates IFNγ production at late stages of T cell activation in an ARE-dependent manner. **(A)** Schematic representation of the experimental setup of WT or KO OT-I effector T cell co-cultures with B16-OVA melanoma cells in an 6:1 effector : target ratio. **(B)** Frequency of IFNγ producing cells and IFNγ expression levels for indicated time points. **(C)** Immunoblot of ZFP36L2 and RhoGDI on lysates of WT and KO OT-I T cells transduced with an GFP empty vector (EV) construct (left and middle lane) and KO OT-I T cells transduced with an ZFP36L2-IRES-GFP construct (right lane). **(D)** IFNγ production in WT, KO and ZFP36L2 reconstituted KO T cells after co-culture with B16-OVA cells for indicated time points. (B, D) Brefeldin A was added for the last 2 h of activation. **(E)** Representative histograms of GFP of T cells that were co-transduced with Katushka-EV or Katushka ZFP36L2, and full-length WT-*Ifng* 3’UTR or AREmut *Ifng* 3’UTR fused to GFP. Numbers in the upper corners indicate GFP gMFI levels of indicated constructs. **(F)** Quantification of GFP gMFI levels in T cells transduced with indicated GFP fusion constructs after co-culture with B16-OVA cells for 48h (n=3 mice/group). **(G)** Relative contribution of ZFP36L2 to regulate GFP gMFI levels of full-length 3’UTR *Ifng* GFP reporter constructs compared to ARE-mutated 3’UTR *Ifng* GFP reporter constructs (n=3 mice/group) in WT and KO T cells after co-culture with B16-OVA for 48h. For calculation see methods. Data were analyzed by two-sided, unpaired Student’s *t*-test, mean ± s.d. **(B)**; *p<0.05, **p<0.01, or by one-way ANOVA with Tukey multiple comparison correction, mean ± s.d. (**D,F,G**); *p<0.05, **p<0.01, ***p<0.001, ****p<0.0001.

We first re-introduced ZFP36L2 expression into ZFP36L2^CKO^ T cells by retroviral transduction to determine whether the effect of ZFP36L2 on IFNγ was direct, or due to indirect effects (**Fig. 5C**). For analysis, we gated on GFP positive T cells **(Supp. Fig. 5A)**. Retroviral transduction with ZFP36L2-IRES-GFP, but not with the empty-GFP construct, reversed the increased IFNγ production in ZFP36L2^CKO^ OT-I T cells (**Fig. 5D**). Re-expression of ZFP36L2 restored the regulation of IFNγ production on percentage and per cell cytokine production, and this effect was most pronounced at 48h of co-culture with B16-OVA cells (**Fig. 5D**). Thus, the observed effects on cytokine production in ZFP36L2^CKO^ were directly instructed by ZFP36L2.

We previously showed that ZFP36L2 directly interacts with *Ifng* mRNA in quiescent memory T cells through AREs within the *Ifng* 3’UTR (35). Upon T cell activation, however, ZFP36L2 rapidly releases the preformed *Ifng* mRNA (35). We therefore questioned whether ZFP36L2 directly interacts with *Ifng* mRNA during later time points of T cell activation, when the effect of ZFP36L2 deficiency was most prominent. WT and ZFP36L2^CKO^ OT-I T cells were transduced with GFP 3’UTR reporter constructs containing the wildtype sequence of the *Ifng* 3’UTR (WT) or the *Ifng* 3’UTR with the 5 ARE sequences mutated (AREmut) (35). As a negative control, we used the perforin 3’UTR, and GFP empty control vector. To reintroduce ZFP36L2 expression, we co-transduced ZFP36L2^CKO^ OT-I T cells with ZFP36L2-IRES-Katushka. WT cells were transduced with empty-Katushka constructs. For analysis, GFP/Katushka double positive T cells were used (**Supp. Fig. 5B**).

The GFP expression of cells transduced with GFP empty control, or perforin 3’UTR was stable throughout in WT, ZFP36L2^CKO^, or ZFP36L2 reconstituted ZFP36L2^CKO^ cells (**Sup. Fig. 5C-E)**. Conversely, and comparable to the endogenous IFNγ protein (**Fig. 5D**), GFP reporter expression was suppressed by the WT *Ifng* 3’UTR in WT OT-I T cells, but much less so in ZFP36L2^CKO^ ko (**Fig. 5F**). Reconstitution of ZFP36L2 in ZFP36L2^CKO^ cells restored the 3’UTR mediated regulation of GFP reporter expression (**Fig 5F**, right panel). Importantly, the *Ifng* 3’UTR-mediated regulation of GFP expression was lost when AREs were mutated, irrespective of the status of ZFP36L2 within T cells (**Fig 5F**). Thus, also during late T cell activation, ZFP36L2 regulates IFNγ through AREs in the 3’UTR.

To determine the relative contribution of ZFP36L2 on GFP protein expression, we used the ratio between GFP WT *Ifng* 3’UTR and the GFP ARE-mut *Ifng* 3’UTR. Whereas ZFP36L2 displayed only a slight inhibitory effect on GFP expression at 6h co-culture with B16-OVA cells, its effect on GFP expression increased at day 1, and even more so at day 2 of activation upon coculture with B16-OVA, explaining almost 45% of the ZFP36L2 mediated regulation of *Ifng* 3’UTR (**Fig. 5G**). Thus, ZFP36l2 regulates IFNγ expression in continuously activated effector T cells through AREs located within the *Ifng* 3’UTR.

### ZFP36L2-mediated regulation of IFN**γ** production occurs independently of activation-induced phosphorylation

We then questioned how the activity of ZFP36L2 is regulated. RBP activity can be modulated by post-translational modifications (48, 49), including phosphorylation (50). To determine whether ZFP36L2 becomes phosphorylated upon T cell activation, we used a Phos-tag SDS-page that separates phosphorylated from unphosphorylated proteins. Indeed, within 5 minutes of T cell activation with the N4 peptide, phospho-ERK1/2 appeared (**Sup. Fig. 6A**). Using the flag-tagged WT ZFP36L2 in ZFP36L2^CKO^ OT-I T cells, we found that within 1h of T cel activation, ZFP36L2 was phosphorylated (**Sup. Fig. 6A**). Published phospho-proteomics studies on murine T cells identified three phosphorylation sites (Ser57, Ser59, and Ser127) of ZFP36L2, of which Ser127 showed increased phosphorylation during both short term (i.e. 1-2h) and long term (16h) T cell activation (**Sup. Fig. 6B**). To determine whether phosphorylation of the indicated serine residues defines the function of ZFP36L2, we generated single and triple mutants of these serine residues to alanine. Flow cytometry analysis with the α-flag antibody revealed that all ZFP36L2 mutants were expressed in ZFP36L2^CKO^ OT-I T cells (**Sup. Fig. 6C**). To determine the functional consequence of ZFP36L2 phosphorylation, we co-cultured reconstituted ZFP36L2^CKO^ OT-I T cells with B16-OVA tumour cells. After day 1 of co-culture with B16-OVA cells, only the ZFP36L2 triple mutant, but none of the other ZFP3L2 mutants showed a slight attenuation of IFNγ production, a feature that was lost at 48h of activation (**Sup. Fig. 6D, E**). We therefore conclude that phosphorylation of ZFP36L2 on these serine residues are not the main drivers of ZFP36L2 function, at least not when T cells are continuously activated.

### ZFP36L2 dampens IFN**γ** production in chronically stimulated T cells by destabilizing ***Ifng* mRNA**

Lastly, we studied the mode of action that ZFP36L2 employs to attenuate the production of IFNγ upon continuous cytokine production. ZFP36L2^CKO^ T cells displayed a 10-fold increase of *Ifng* mRNA levels compared to WT T cells when continuously exposed (48h) to B16-OVA cells (**Fig. 6A**). Again, this effect on *Ifng* mRNA levels by ZFP36L2 deficiency could be reversed by reconstituting T cells with WT ZFP36L2 (**Fig. 6A**). In line with this finding, *Ifng* mRNA was stabilized in ZFP36L2^CKO^ OT-I T cells, as defined by measuring the mRNA decay upon blocking *de novo* transcription with Actinomycin D (ActD, **Fig 6B**). Importantly, we also observed slightly higher *Ifng* mRNA levels in ZFP36L2^CKO^ OT-I TILs compared to WT OT-I TILs isolated from B16-OVA tumours (**Fig. 6C**, left panel). The *Ifng* mRNA was also stabilized in chronically stimulated tumour-derived ZFP36L2^CKO^ OT-I T cells, which we observed both when looking at relative *Ifng* mRNA levels, and when measuring the *Ifng* mRNA decay (**Fig. 6C**, right panel). Thus, as opposed to the translational block that ZFP36L2 confers on cytokine mRNA in quiescent memory T cells (35), in effector T cells ZFP36L2 destabilizes *Ifng* mRNA, and it does so primarily at late time points of T cell activation.

**Figure 6.**
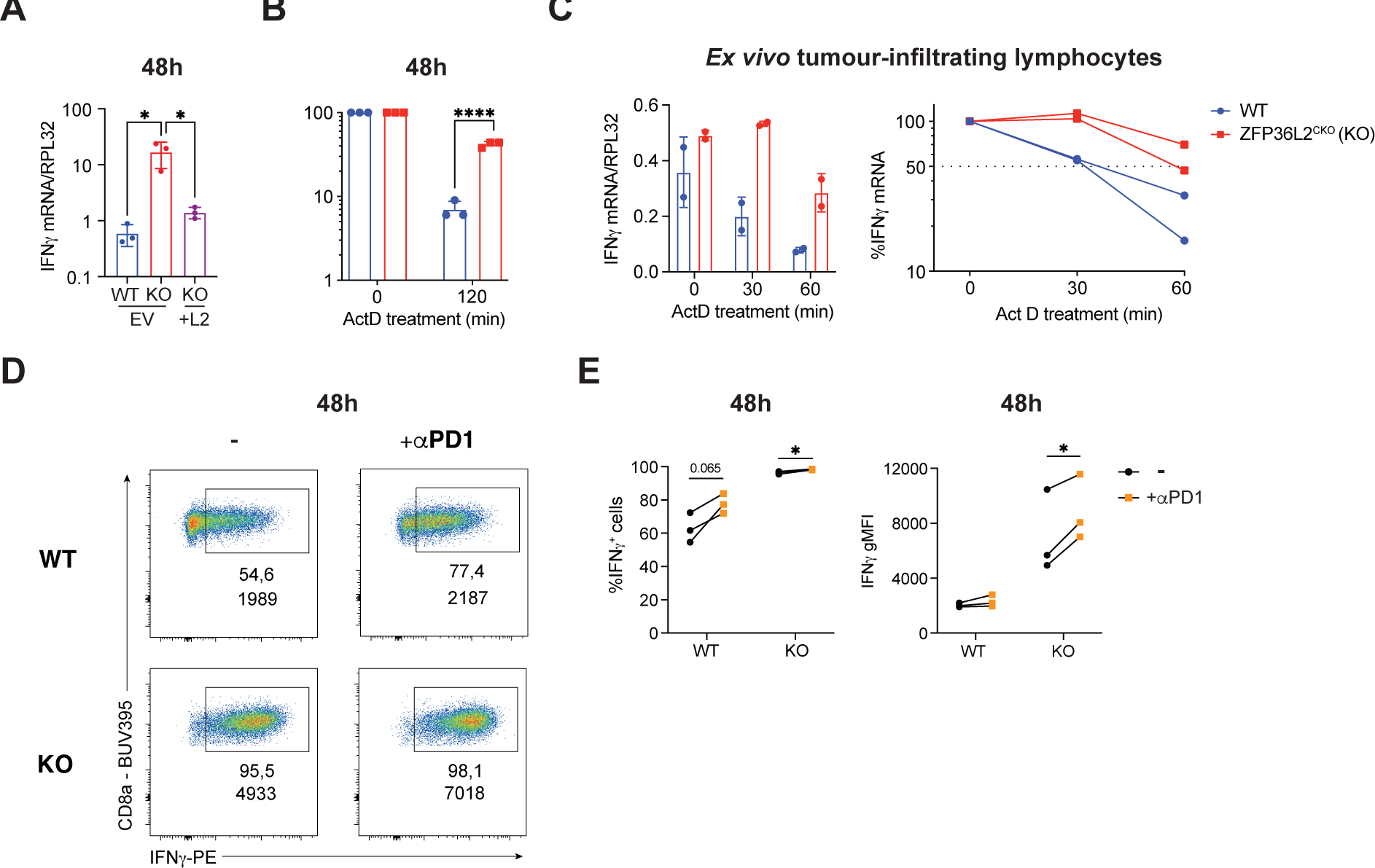
ZFP36L2 destabilizes *Ifng* mRNA during prolonged T cell activation and acts independently of PD-1 signalling. **(A)** *Ifng* mRNA expression in sorted WT, KO and, KO reconstituted with WT ZFP36L2, T cells, 48h after continuous co-culture with B16-OVA. **(B)** *Ifng* mRNA decay in WT and KO after 48h of continuous co-culture with B16-OVA cells. T cells were treated with 1μg/ml Actinomycin D (ActD) for indicated time points. **(C)** *Ifng* mRNA levels in WT and KO TILs that were sorted and rested for 30 minutes. Total RNA was isolated and mRNA levels were determined at 0h. RNA decay was measured by ActD treatment for indicated time points. **(D-E)** Representative flow cytometry plots (D), and quantification (n=3 mice/group) (E) of WT and KO T cells that were co-cultured for 48h with B16-OVA tumour cells in the presence or absence of PD-1 blocking antibody. Numbers indicate the frequency of IFNγ producing cells and the IFNγ gMFI levels. Data were analyzed by one-way ANOVA with Tukey multiple comparison correction, mean ± s.d. (**A**), by two-sided, unpaired Student’s *t*-test, mean ± s.d. (**B,C**) or by two-sided paired Student’s *t*-test, mean ± s.d. (**E**); *p<0.05. *p<0.05; **p<0.01, ***p<0.001, ****p<0.0001.

We next reasoned that if the mode of action of ZFP36L2 in effector T cells is primarily through mRNA destabilization, the production of IFNγ might be even further increased by boosting translation. We previously showed that αPD1 blockade increased the IFNγ protein production during continuous antigen exposure without altering the *Ifng* mRNA levels or stability (47). Indeed, including αPD1 blocking antibodies to WT and ZFP36L2^CKO^ OT-I T cells in the B16-OVA co-culture system revealed that the IFNγ protein production was not only increased in WT OT-I T cells (**Fig. 6D, E**), but also that αPD1 blockade further increased the production of IFNγ in ZFP36L2^CKO^ T cells (**Fig. 6D, E**). Thus, combined targeting of ZFP36L2 and PD1 blockade synergistically increases the production of IFNγ in CD8^+^ T cells in response to target cells.

## DISCUSSION

In this report, we determined the role of ZFP36L2 during T cell differentiation and activation. Whereas ZFP36L2 is redundant during T cell differentiation, at later stages of T cell activation it dampens IFNγ protein production by destabilizing *Ifng* mRNA.

Intriguingly, we found that the mode of action of ZFP36L2 is context-dependent. We previously reported that ZFP36L2 prevents translation of pre-formed *Ifng* mRNA in quiescent memory T cells (35). Here, we show that ZFP36L2 does not dampen cytokine production early on, but only during later stages of T cell activation. This effect is most profound for IFNγ during continuous stimulation, including TILs. This late effect on cytokine production can be partially explained by changes in ZFP36L2 protein expression: whereas ZFP36L2 protein levels remain stable during the first 12 hours of activation of naive CD8^+^ T cells, they increased about 2-fold after 18 hours of activation (53). However, because ZFP36L2 is expressed throughout, we argue that other RBPs may outcompete, or control its RNA binding activity and regulatory function at early T cell activation. Recently, several RBPs that regulate cytokine production during early T cell activation were identified, including ZFP36 and ZFP36L1 (23, 34). It is therefore conceivable that ZFP36L1 or other ARE-binding RBPs outcompete ZFP36L2 binding during the early phase of T cell activation.

It remains to be established how ZFP36L2 acquires the ability to destabilize mRNA during late T cell effector stages, but not in quiescent memory T cells. It is conceivable that ZFP36L2-intrinsic changes such as post-translational modifications (PTMs) may define its activity (49, 54, 55). Even though we found no evidence that the previously identified serine residues S57, S59 and S127 (51, 52) are involved, PTM prediction software (musite.net, confidence score > 0.5) identified 161 putative PTM sites for ZFP36L2, including 38 phosphorylation sites (6 threonine and 32 serine residues) and 1 methylation site (1 lysine residue). These putative PTM sites include amino acids present in the conserved C-terminus of ZFP36L2, which directly interacts with the CCR4-NOT mRNA degradation complex. It therefore cannot be excluded that PTM defines the function and/or RNA-binding capacity of ZFP36L2. Alternatively, changes in binding partners for ZFP36L2 during different T cell states could potentially define its function. Furthermore, because the RBP expression landscape alters upon T cell differentiation and activation (21, 57), the availability of RBP interaction partners for ZFP36L2 may change. Irrespective of the underlying mechanism, we report here that the regulatory mechanisms that ZFP36L2 employs are context-dependent.

IFNγ is required for T cells to control tumour growth (17, 58, 59). Surprisingly, even though ZFP36L2^CKO^ TILs produced more IFNγ, it was not enough for tumour control. ZFP36L2 deficiency increased only ∼50% of IFNγ production when compared to germline deletion of AREs within the *Ifng* 3’UTR (47). Because IFNγ can be sequestered by the extracellular matrix (60), the levels that ZFP36L2^CKO^ TILs reach may not be sufficient for blocking the tumour outgrowth. Furthermore, other T cell effector molecules such as TNF, IL2, CD107 and Granzyme B were unaffected in ZFP36L2^CKO^ TILs, and such broader alterations may be required when the increase of IFNγ is limited (34, 61–63). Additionally, the antitumour effects of IFNγ may be outcompeted by the pro-tumourigenic function of IFNγ, including the induction of PDL1 and IDO1 (64). In line with this hypothesis, PD-1 blockade was sufficient to boost the IFNγ production in ZFP36L2^CKO^ T cells.

In conclusion, we identified a novel context-dependent role for ZFP36L2 in regulating IFNγ production. Considering the important role of IFNγ in controlling tumour growth, we postulate that combinatorial targeting of post-transcriptional regulatory nodes that affect IFNγ production, like ZFP36L2 and PD-1, can further potentiate the effectiveness of T cell therapies against cancer.

## ACKNOWLEDGMENTS

We thank the animal caretakers of the NKI (Netherlands Cancer Institute) and the Sanquin Flow cytometry facility; S. E. Bell, M. Hoogenboezem, B. Popovic, A. Laurent, T. Verkerk for technical help; and the Turner lab and B. Popovic for critical reading of the manuscript. Mice were generated under Biotechnology and Biological Sciences Research Council grant BBS/E/B/000C0428. This research was supported by the European Research Council consolidator grant PRINTERS 817533.

## SUPPLEMENTARY FIGURE LEGENDS

**Supplementary Figure 1.**
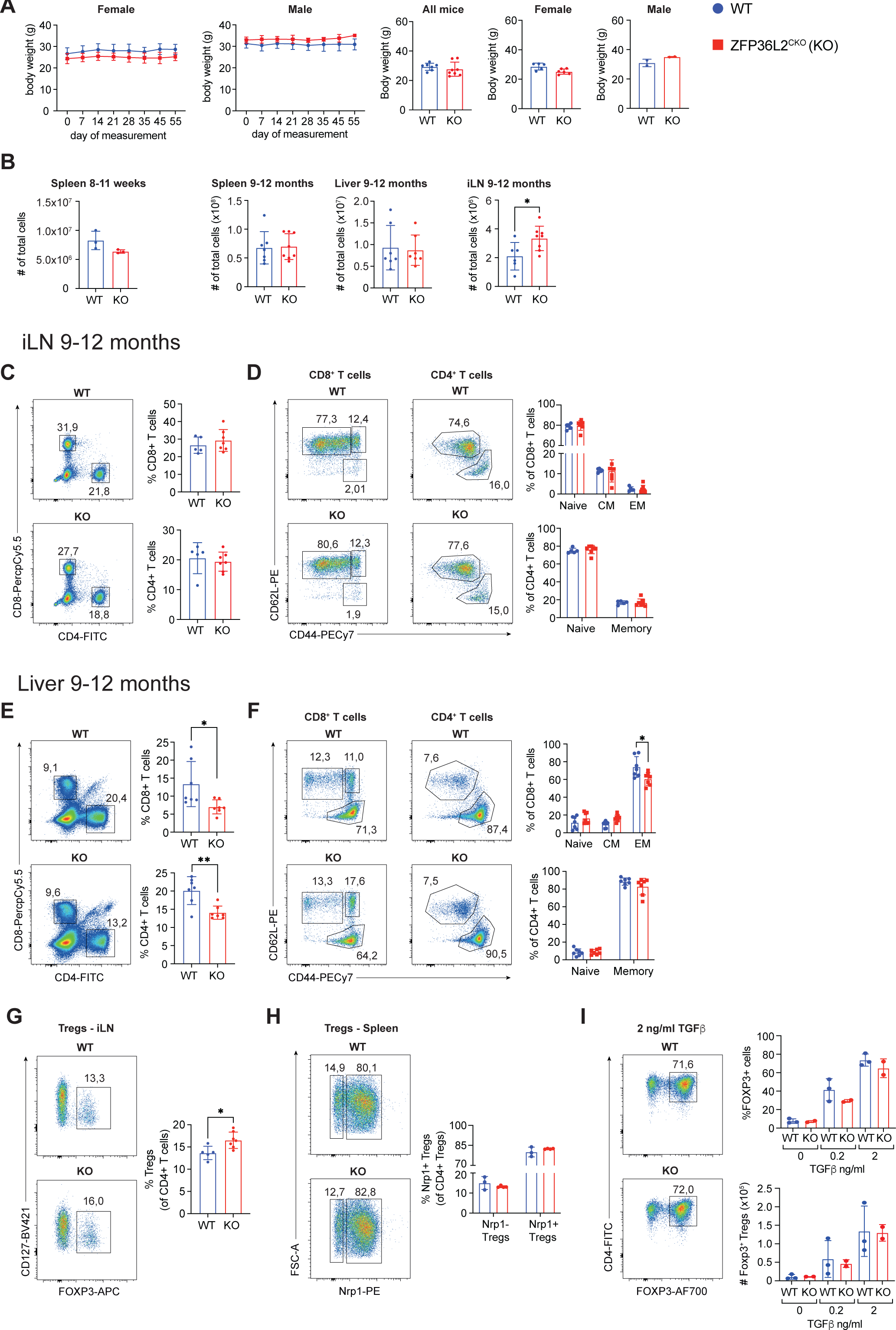
Phenotyping of WT and ZFP36L2^cko^ littermates. **(A)** Body weight of 9-12 months old female (n=5 for WT and n=6 for KO) and male (n=2/group) mice measured over a period of 55 days. (Right panel) Body weight at end point (day 55) of aging for all mice (n=7 for WT mice and n=8 for KO mice), female mice (n=5 WT and n=6 KO mice) and male mice (n=2 mice/group). **(B)** Absolute cell numbers isolated from spleens of young mice (8-11 weeks; left panel, n=3 mice/group) and of 9-12-month-old mice from spleen (n=7 for WT mice, n=8 for KO mice), liver (n=7 mice/group) and iLN (right panel, n=6 for WT mice, n=8 for KO mice). **(C-F)** Representative flow cytometry plots and frequencies of CD8^+^ and CD4^+^ T cell populations (C, D) and their respective T cell differentiation phenotype (D, F) including naive (CD62L^+^ CD44^-^), central memory (CD62L^+^ CD44^+^) and effector memory (CD62L^-^ CD44^+^) CD8^+^ T cells, and naïve (CD62L^+^ CD44^low^) and memory (CD62L^-^ CD44^hi^) CD4^+^ T cells isolated from indicated organ (n=7 mice/group). **(G)** Frequencies of CD4^+^ Tregs from iLN (CD127^low^ Foxp3^+^; n=5 for WT mice and n=7 for KO mice). **(H)** Representative flow cytometry plots and quantification of Nrp1^-^ and Nrp1^+^ splenic Tregs (8-1 weeks old). **(I)** Representative flow cytometry plots and quantification of frequency and absolute number of *in vitro* generated iTregs for WT and KO cells (n=3 WT and n=2 KO mice). Data were analyzed by one-way ANOVA with Tukey multiple comparison correction, mean ± s.d. (**D,F,H,I**); or by two-sided, unpaired Student’s *t*-test, mean ± s.d. (**A-C,E,G**). *p<0.05, **p<0.01.

**Supplementary Figure 2.**
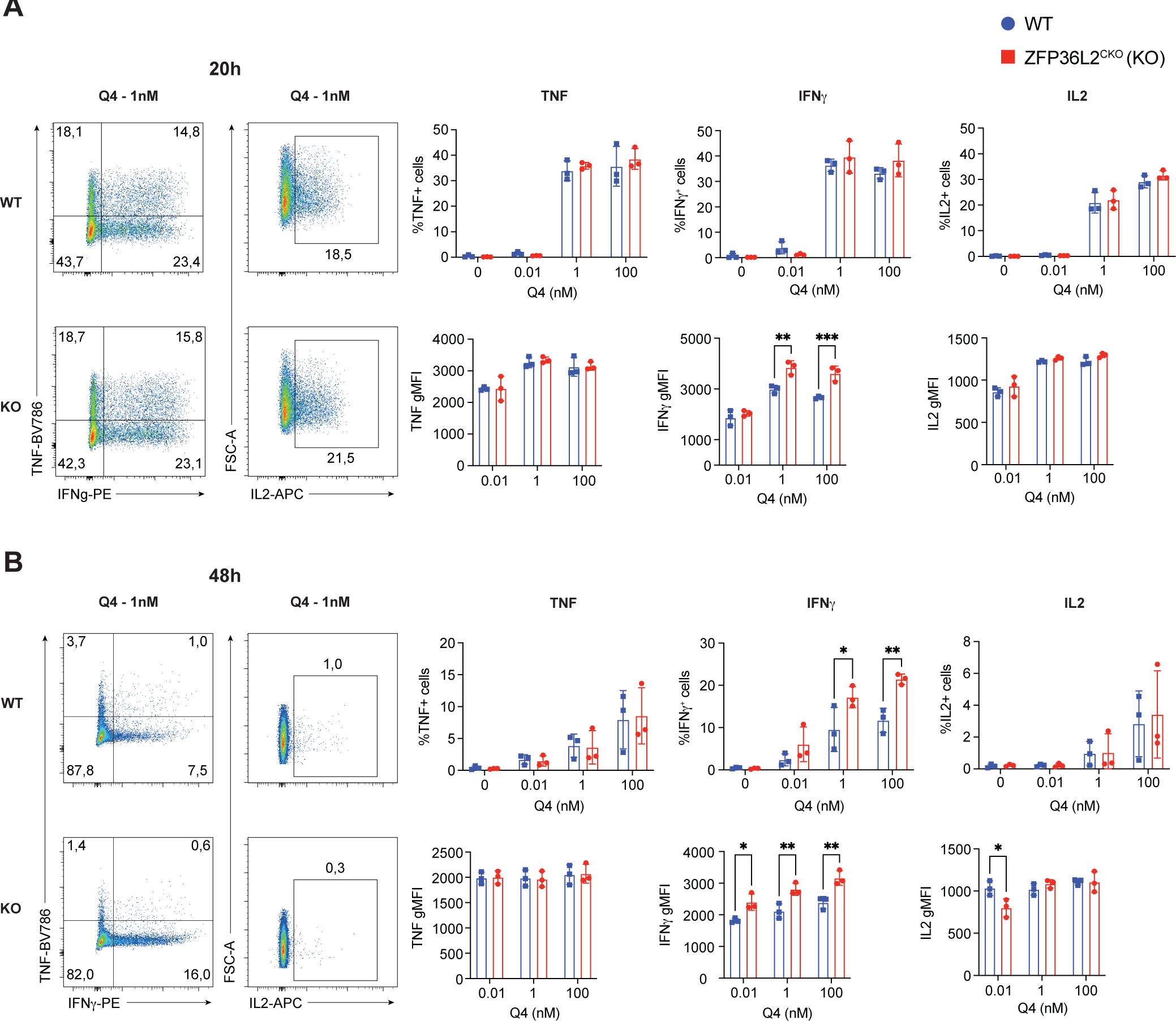
ZFP36L2 negatively regulates IFNγ production in naive CD8 T cells during priming. **(A,B)** Cytokine production of naive WT and KO OT-I CD8^+^ T cells that were primed for 20 h and 48h with bone marrow-derived DCs loaded with indicated concentrations of the low affinity SIIQFEKL (Q4) peptide. Brefeldin A was added for the last 2 hours of each time point. (n=3 mice/group). Percentage (top right) and cytokine production per cell (bottom right) as defined by the Geo-mean fluorescence intensity (gMFI). Data were analyzed by two-way ANOVA with Šidák multiple comparison correction, mean ± s.d. ; *p<0.05; **p<0.01.

**Supplementary Figure 3.**
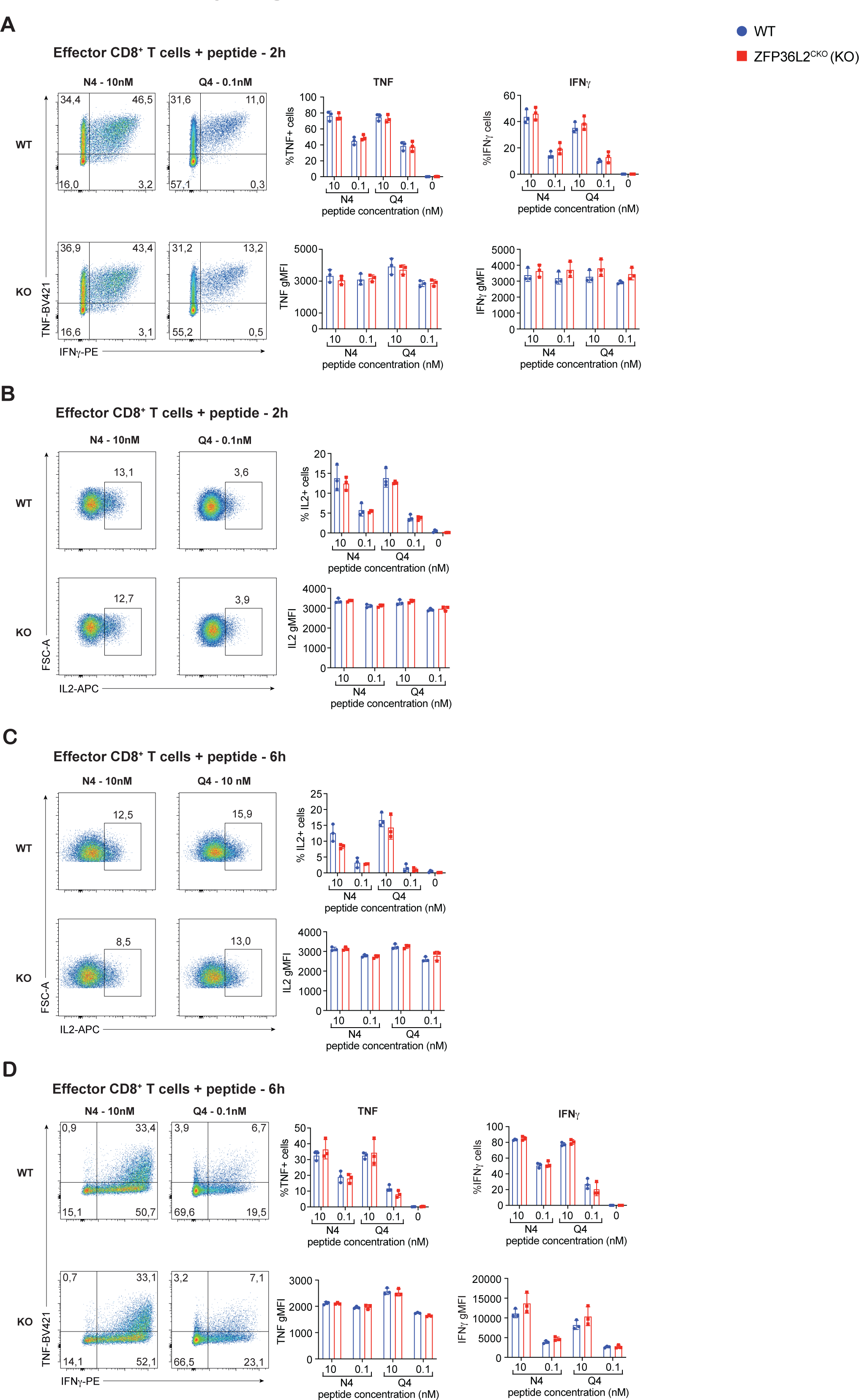
ZFP36L2 deficiency does not affect cytokine production upon short term T cell activation of effector and virtual memory T cells. **(A-D)** Cytokine production of *in vitro* rested WT and KO OT-I T cells after activation with different concentrations of the N4 or Q4 peptide for 2h (A,B) and 6h (C,D). Brefeldin A was added for the last 2 h of activation. (n=3 mice/group). Data were analyzed by two-way ANOVA with Šidák multiple comparison correction, mean ± s.d.

**Supplementary Figure 4.**
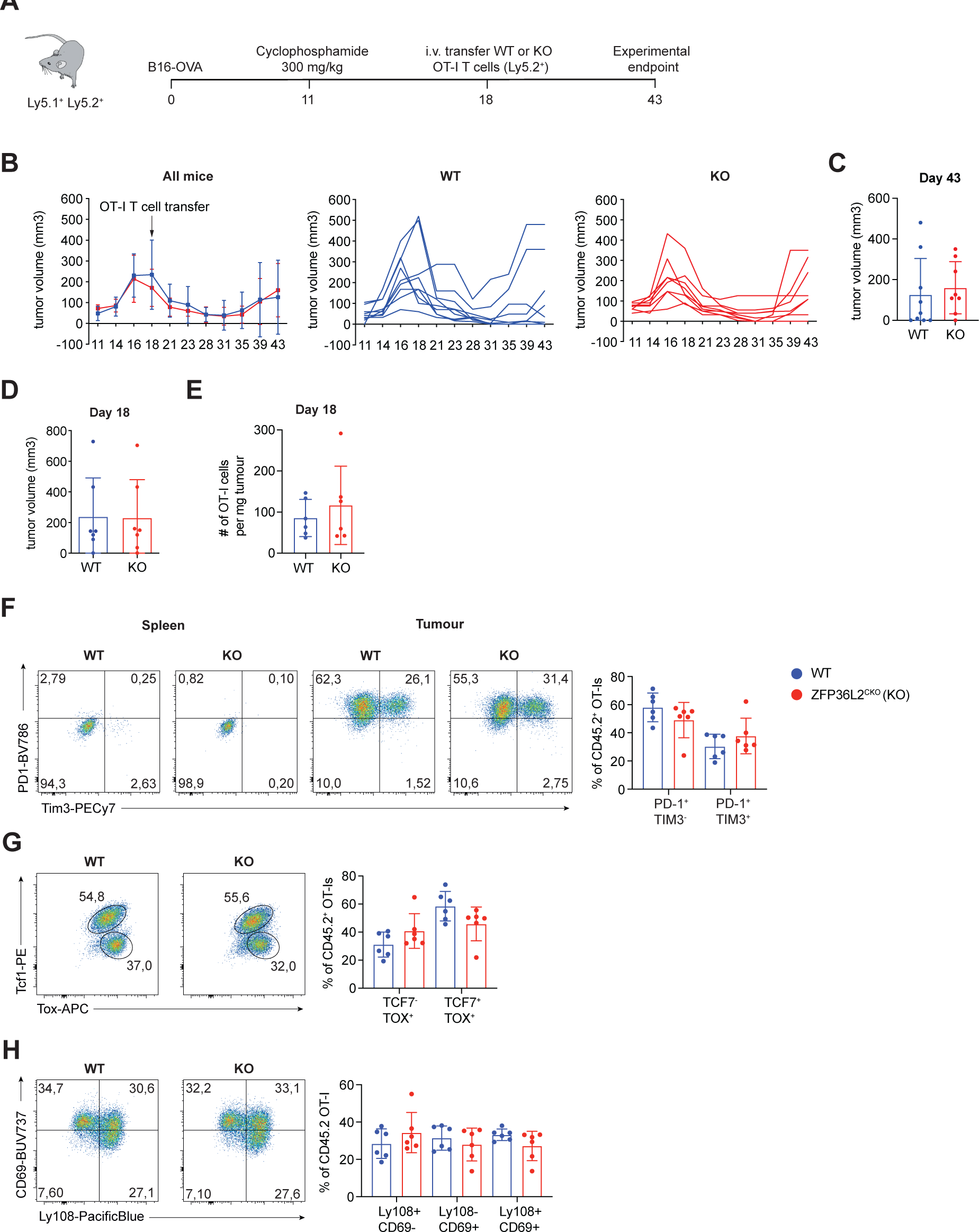
ZFP36L2 deficiency in TILs does not influence T cell differentiation, or tumour growth. **(A)** Schematic overview of *in vivo* B16-OVA tumour growth experiment. Ly5.1^+^ Ly5.2^+^ recipient mice were engrafted with 1 x 10^6^ B16-OVA cells subcutaneously. **(B)** Tumour growth curve of all mice combined and of individual mice that received adoptive T cell therapy of WT or KO OT-I T cells (n=9 for WT mice, n=8 for KO mice). **(C)** Tumour size at day 43 (25 days after T cell transfer). n=9 for WT mice, n=8 for KO mice. **(D)** Tumour volume of B16-OVA bearing mice at day 18 that had received WT or KO OT-I T cells (n=7 mice/group). **(E)** Absolute number of WT or KO OT-I T cells per gram of tumour (n=6 mice/group). **(F)** Representative flow cytometry plots and frequencies of PD-1^+^ Tim3^-^ and PD-1^+^ Tim3^+^ WT and KO OT-I splenocytes and TILs (n=6 mice/group). **(G,H)** Representative flow cytometry plots and percentages of Tcf1 and Tox expression (G), and CD69 and Ly108 expression (H) of WT and KO TILs (n=6 mice/group). Data were analyzed by two-way ANOVA with Šidák multiple comparison correction, mean ± s.d. (**F,G,H**), or by two-sided, unpaired Student’s *t*-test, mean ± s.d. (**C,D,E)**.

**Supplementary Figure 5.**
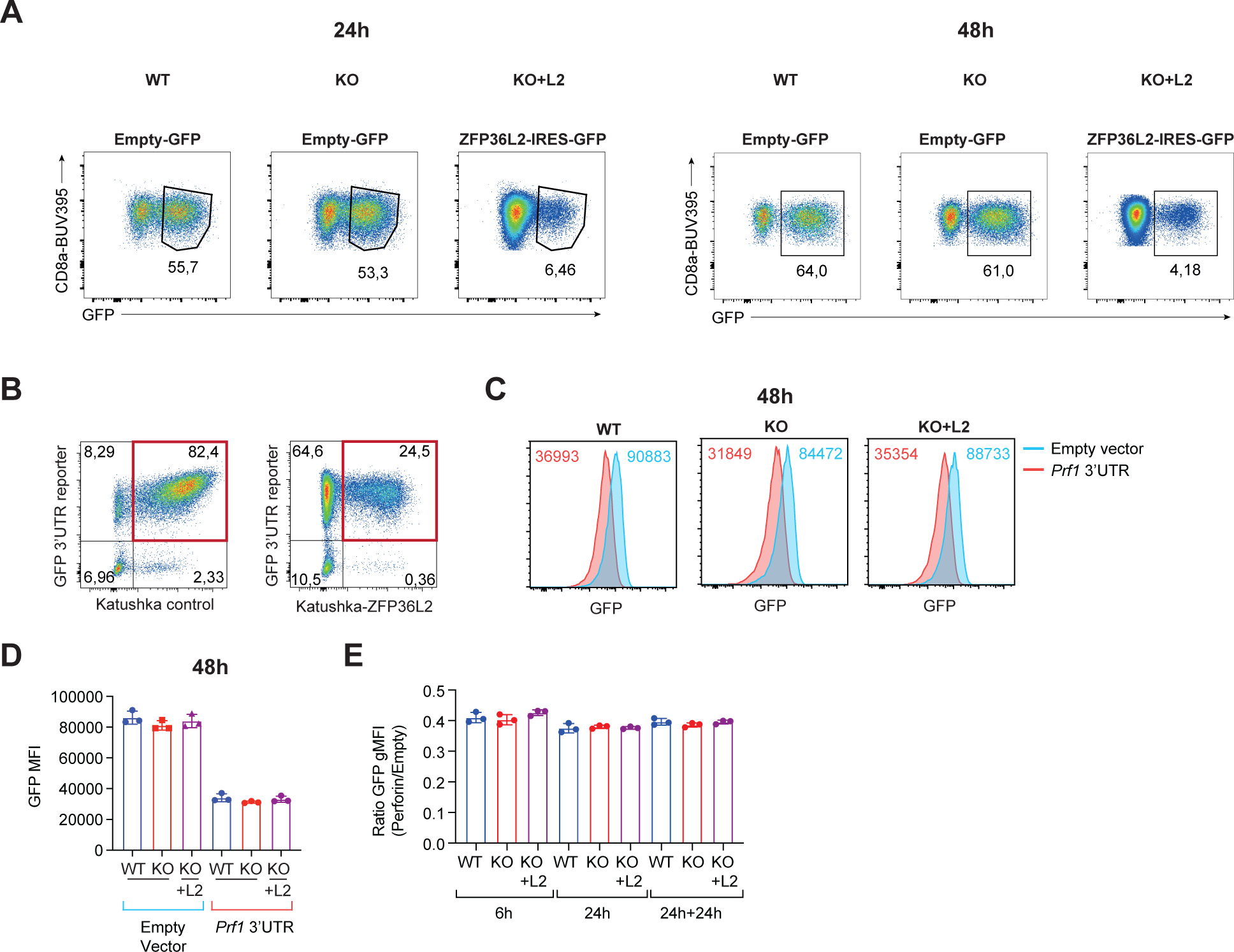
Early cytokine expression and the 3’UTR mediated-expression of Perforin is not regulated by ZFP36L2. **(A)** Gating strategy for OT-I T cells transduced with an empty-GFP control construct or ZFP36L2-IRES-GFP construct. **(B)** Gating strategy for OT-I T cells co-transduced with GFP *Ifng* 3’UTR reporter and Katushka empty vector control (left panel) or ZFP36L2-IRES-Katushka (right panel) constructs. **(C)** Representative histograms of GFP expression among T cells transduced with GFP control or GFP *Prf1* 3’UTR fusion constructs. Numbers in the upper left corner display GFP gMFI levels of indicated constructs. **(D)** Quantification of GFP gMFI levels (n=3 per group) among T cells transduced with GFP control or GFP *Prf1* 3’UTR fusion constructs. **(E)** Relative GFP gMFI levels of full-length 3’UTR *Prf1* GFP reporter constructs compared to GFP control reporter constructs (n=3 per group). Data were analyzed by one-way ANOVA with Tukey multiple comparison correction, mean ± s.d. **(D,E);** *p<0.05; **p<0.01, ***p<0.001, ****p<0.0001.

**Supplementary Figure 6.**
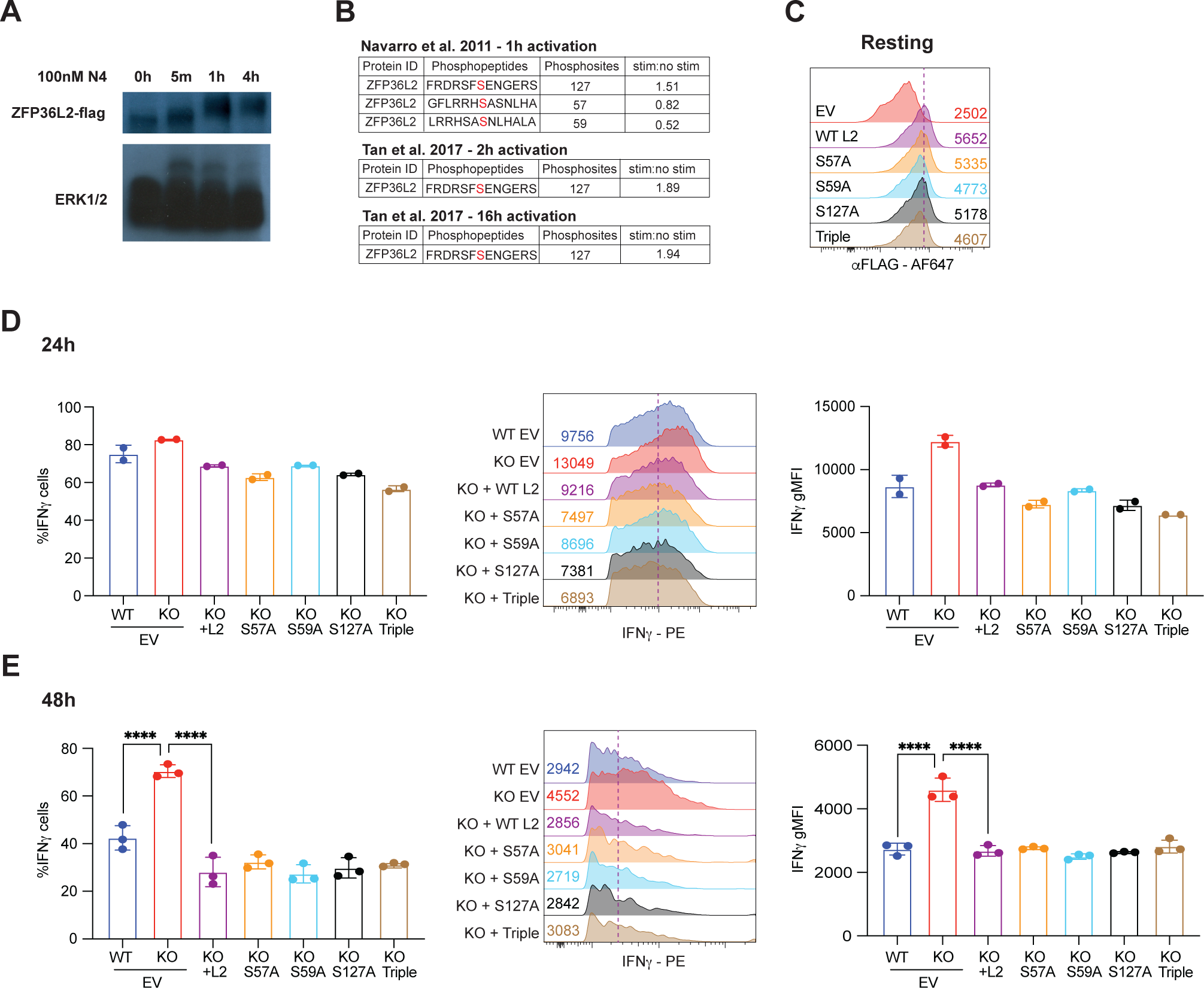
Phosphorylation of ZFP36L2 at S57, S59 and S127 is not required for its functionality. **(A)** Phostag blot of ZFP36L2-flag and ERK1/2 expression in OT-I T cells after activation with 100nM N4 peptide for indicated timepoints. **(B)** ZFP36L2 phosphopeptides that were extracted from phosphoproteomics datasets from Navarro *et al.* (51) and Tan *et al.* (52) from mouse T cells that were activated for indicated time points. **(C)** Expression of ZFP36L2 mutants in reconstituted resting KO OT-I T cells shown by representative histograms. GFP gMFI levels are indicated on the right. **(D-E)** IFNγ production of WT and KO that were transduced with an empty GFP control construct, and KO OT-I T cells that were reconstituted with WT or mutant ZFP36L2 constructs and that were cocultured with B16-OVA cells for 24h (F, n=2mice/group) and for 48h (G, n=3mice/group). Data were analyzed by one-way ANOVA with Tukey multiple comparison correction, mean ± s.d. (**D,E**); *p<0.05; **p<0.01, ***p<0.001, ****p<0.0001.

**Supplemental Table I.**
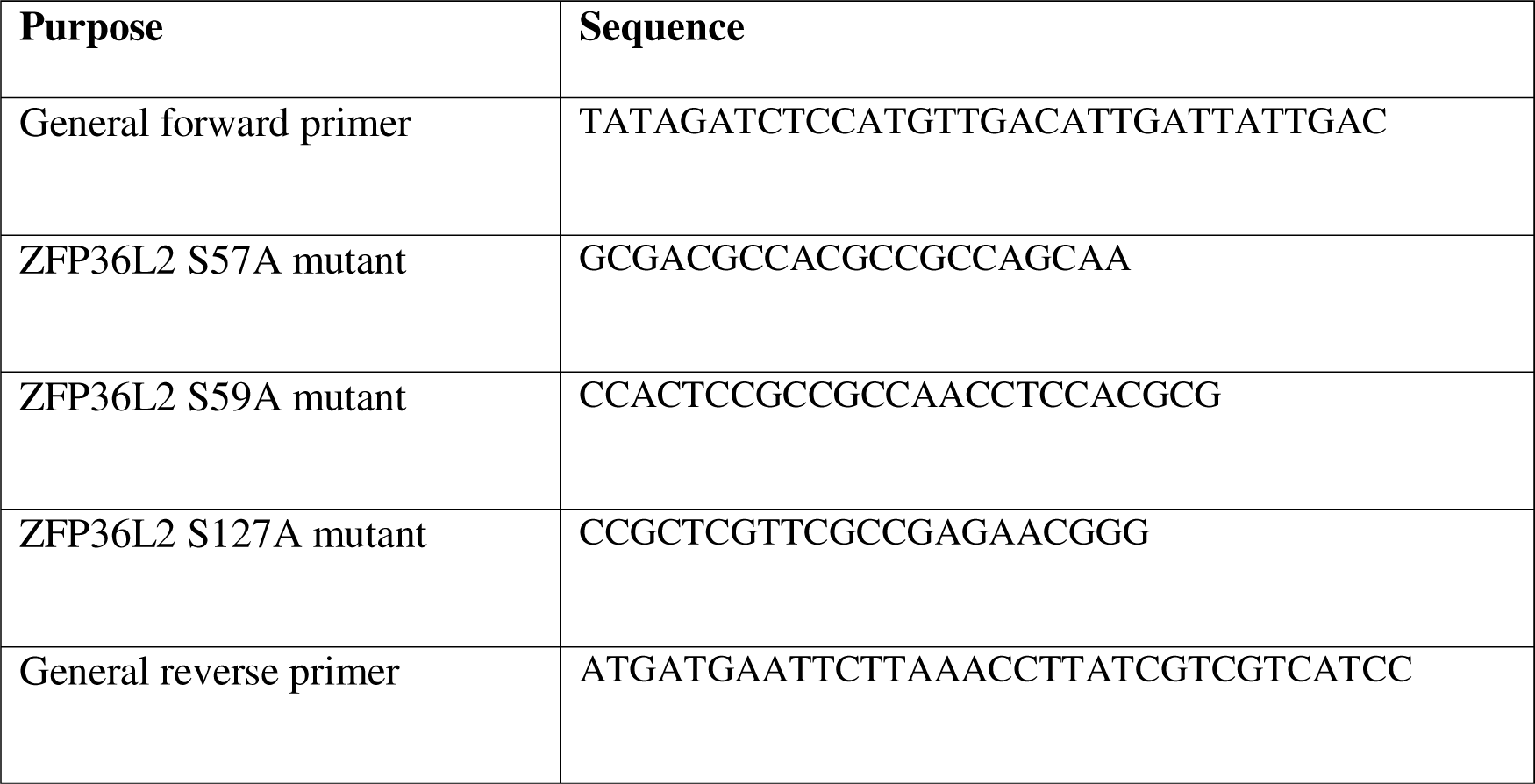

## Material and Methods

### Mice and cell culture

C57BL/6j/Ly5.1^+^Ly5.2^+^ mice and Zfp36l2;CD4cre mice were bred and housed in the animal facility of the Netherlands Cancer Institute (NKI). Zfp36l2;CD4cre (ZFP36L2^CKO^) mice were crossed with C57BL/6j/Ly5.2 OT-I mice to obtain ZFP36L2^CKO^ and WT littermate OT-I T cells (65). Animals were genotyped by Transnetyx. Animals were housed in individually ventilated cage systems under specific pathogen-free conditions. All experiments were performed in accordance with institutional and national guidelines and approved by the experimental animal committee of the NKI. Mice were used at age 6-12 weeks of age, unless specified otherwise. B16-OVA cells (44) and MEC.B7.SigOVA cells (38) were cultured at 37°C and 5% CO_2_ in IMDM (GIBCO-BRL) containing L-glutamine and 25mM HEPES, supplemented with 10% FCS, 15μM 2-mercaptopethanol, 2mM L-glutamine, 100 U/ml penicillin G sodium and 100 μg/ml streptomycin sulfate (complete medium). PlatE cells (66) were cultured at 37°C and 5% CO_2_ in DMEM containing 4.5g/l D-glucose, L-glutamine and 25mM HEPES supplemented with 2mM L-glutamine and 100 U/ml penicillin G sodium and 100 μg/ml streptomycin sulfate.

### Plasmids and mutagenesis

Gene-specific 3’UTR GFP fusion reporter constructs were generated as previously described (35). C-terminal flag-tagged ZFP36L2 (Origene) was cloned into the pMIG-W vector (67) using the BgLII and EcoRI restriction sites. ZFP36L2 S57A, S59A and S127A mutants were generated using two forwards primers; one primer at the N-terminal start site of the CDS and one primer with up to two non-complementary nucleotide changes in the centre of the sequence, and a reverse primer that anneals at the C-terminal site of the CDS (**Supp. Table 1**). PCR reaction was performed with high-fidelity Q5 polymerase (NEB) according to the manufacturer’s protocol. Amplicons were inserted into the pMIG-W backbone. ZFP36L2 Katushka and control vectors were generated by exchanging GFP with Katushka in pMIG-W and pMIG-W-ZFP36L2 using NcoI and PacI. Sequences were confirmed by sequencing analysis.

### T cell isolation and flow cytometry

Mouse organs were harvested and briefly kept in cold PBS on ice until processed. Mouse organs were mashed and passed through a 70μm cell strainer (Corning) and washed with FCS-containing media. Lymphocytes from liver cell-suspensions were enriched by 40-80% percoll (Sigma Aldrich) gradient (860 RCF, 20 min, 21°C, no brake). The interface was washed with PBS (Sigma Aldrich). Cells were washed with FACS buffer (PBS, 1% FCS and 2mM EDTA) and stained for extracellular molecules with the following antibodies: anti-CD8a (53-6.7, BD), anti-CD4 (GK1.4, Thermo), anti-CD25 (PC61.5, eBioscience), anti-CD62L (MEL-14, Biolegend), anti-CD44 (IM7, Biolegend), anti-CD127 (A7R34, Biolegend), anti-NRP-1 (3E12, Biolegend), anti-CD107a (1D4B, eBioscience), anti-PD1 (J43, BD), anti-TIM3 (RMT3-23), anti-Ly108 (330-AJ, Biolegend), anti-CD69 (H1.2F3, BD Horizon)for 20 min at 4°C. Intracellular molecules were stained with the following antibodies: anti-FOXP3 (FJK-16s, Thermo), anti-TNF (MP6-XT22, Biolegend), anti-IFNγ (XMG1.2, Biolegend), anti-IL-2 (JES6-5H4, Thermo), anti-Granzyme B (GB11, BD), anti-TCF (C63D9, Cell Signaling), anti-TOX (REA473, Miltenyi), anti-FLAG (L5, Biolegend) 25 min at 4°C using the BDCytofix/Cytoperm kit (BD Biosciences) according to the manufacturer’s protocol. Transcription factor staining was performed with eBioscience Foxp3/Transcription factor staining kit (ThermoFisher Scientific) according to the manufacturer’s protocol. For intracellular staining of T cells expressing ZFP36L2-flag-IRES-GFP, fixation was performed using 2% paraformaldehyde for 30 min at 4°C (68). Flow cytometry analysis was performed with FACSymphony (BD Biosciences) or LSRFortessa (BD Biosciences). Data were analyzed with FlowJo (BD Biosciences, version 10).

### T cell activation and culture

ZFP36L2^CKO^ and WT OT-I T cells were purified with the MACS CD8a^+^ T cell isolation kit (Miltenyi Biotec) according to the manufacturer’s protocol. Effector OT-I T cells were generated by activating purified OT-I T cells (1e6 cells/well) with pre-seeded MEC.B7.SigOVA cells (1e5 cells/well) for 20 h. T cells were removed from stimulation and cultured in complete medium supplemented with10ng/ml recombinant mouse IL-7 (Peprotech) for 3-5 days as described (69).

Naive and memory CD8^+^ T cells were FACS-sorted from bulk CD8a^+^ T cells using CD62L and CD44. For iTreg induction, naive CD4^+^ T cells were activated with αCD3/αCD28 beads (1:1 cell:bead ratio, ThermoFisher) in the presence of TGFB (Biolegend) and IL-2 (5ng/ml, Miltenyi) for 4 days. Splenocytes from ZFP36L2^CKO^ mice were activated *ex vivo* with 10ng/ml PMA and 1 μM Ionomycin. Naïve CD8^+^ T cells were activated with bone marrow-derived DCs loaded for 1 h with indicated concentrations of N4 or Q4 OVA_257-264_ peptide versions (Anaspec) as previously described (69). *In vitro* reactivation of OT-I T cells was performed with indicated concentrations of N4 or Q4 OVA_257-264_ peptide. Memory CD8^+^ T cells were activated with αCD3/αCD28 beads (1:1 cell:bead ratio, ThermoFisher). For cytokine measurements, OT-I T cells were incubated with 1 μg/ml brefeldin A (BD Biosciences) for the last 2 hours of activation.

### Retroviral transduction

PlatE cells were plated in 6-well plates (0.5e6/well) and cultured overnight. The following day, media was substituted with 900ul of pre-warmed media. After 30 min, PlatE cells were transfected with 1μg of retroviral vector of interest, together with 1μg pCL-Eco, using GeneJammer in a 2:6 ratio, according to the manufacturer’s protocol (Agilent). 8 h after transfection 1ml of pre-warmed media was added. After 48 hours, virus was harvested, filtered through a 0.45 μm filter and directly used, or snap-frozen using liquid nitrogen and stored at -80°C. Virus was plated on non-tissue culture treated 24-well plates precoated with retronectin (TaKaRa) overnight at 4°C, and centrifuged for 30 min at 4°C at 3599 RCF (5 acceleration ramp, 0 deceleration ramp). Activated OT-I T cells were added, centrifuged for 5 min at 4°C at 1000rpm, and incubated for 24 h at 37°C. Medium was refreshed and cells were rested for 4 days prior to analysis.

### B16-OVA Tumour model

To study T cells infiltrating tumours, 8-12-week-old C57BL/6J/Ly5.1^+^Ly5.2^+^ mice were injected with 1 x 10^6^ B16-OVA cells. On day 7, when tumours were palpable, *in vitro* generated rested effector 1 x 10^6^ ZFP36L2^CKO^ OT1, or WT littermates were injected intravenously. Prior to T cell transfer, dead cells were removed by using Lympholyte (Cedarlane labs). At day 18, animals were culled and spleens and tumours were harvested. Tumours were cut into small pieces and digested with 100μg/ml DNase I (Roche) and 200 U/ml Collagenase IV (Worthington Biochemical) at 37°C for 30 min. Cells were counted and incubated with 1μg/ml brefeldin A and 1μg/ml monensin for 4 h in the presence or absence of 100nM N4 OVA_257-264_ peptide. For tumour growth studies, B16-OVA tumour bearing mice were injected intraperitoneally with 300mg/kg cyclophosphamide at day 11. On day 14, mice received 2 x 10^6^ WT or ZFP36L2^CKO^ OT-I T cells. Tumour growth was tracked until tumours reached ∼1000mm^3^.

For *in vitro* B16-OVA co-culture experiments, in vitro generated effector ZFP36L2^CKO^ and WT OT-I T cells were co-cultured with B16-OVA cells at a 6:1 effector:target ratio for 6h or 24h. 24 h after activation, OT-I T cells were transferred to a fresh plate of pre-seeded B16-OVA cells. PD-1 blocking antibody (29F.1A12, Biolegend) was freshly added (10μg/ml) every day of the co-culture.

To measure the contribution of ZFP36L2 to ARE-mediated regulation, T cells co-transduced with GFP-3’UTR reporter genes and Katushka empty or ZFP36L2-IRES-Katushka containing plasmids were used for the tumour co-culture. To calculate the GFP gMFI ratio the GFP gMFI of WT 3’UTR IFNg samples was divided by the GFP gMFI of ARE-mut 3’UTR IFNg. The relative contribution of ZFP36L2 on regulating the ARE-containing reporter constructs was calculated by dividing the gMFI ratio of WT samples with the relative gMFI ratio between ZFP36L2^CKO^ and WT samples (see formula).

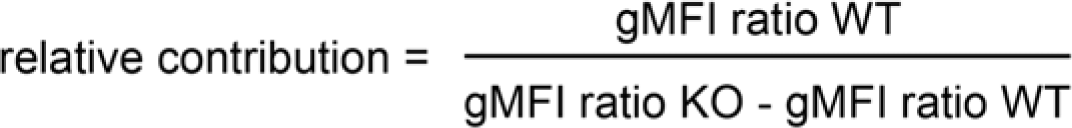

### Phostag and Immunoblotting

Cell lysates (1 x 10^6^ cells per sample) were prepared by standard procedures using RIPA lysis buffer (Thermo) supplemented with protease and phosphatase inhibitors (Thermo). Lysates were run on 4-12% SDS-PAGE (Thermo) or 7.5% Phostag SDS-page (Fujifilm Wako Chemicals). Canonical SDS-PAGE gels were directly transferred onto nitrocellulose membranes (iBlot2, Thermo). Phostag SDS-PAGE gels were first washed with transfer buffers generated and used according to the manufacturer’s protocol and subsequently transferred onto nitrocellulose membranes. Subsequently, membranes were blocked with 5% BSA TBST solution (Fraction V, Sigma). Membranes were stained with α-ZFP36L2 (ab70775, Abcam), α-RhoGDI (MAB9959, Abnova) or α-Flag antibodies (M2, Sigma) followed by α-Rabbit (4050-05, Southern Biotech) or α-Mouse (1031-05, Southern Biotech) HRP-conjugated secondary antibodies.

### RNA extraction and quantitative PCR analysis

Total RNA was extracted from sorted OT-I T cells and extracted using Trizol reagent (Invitrogen). cDNA was synthesized using Superscript III Reverse transcriptase (Invitrogen). Quantitative Real-Time PCR was performed in triplicate with SYBR green on a StepOne Plus (both Applied Biosystems) using previously described primers for *Ifng* and *Rpl32* mRNA (39). Ct values were normalized to RPL32 levels. *Ifng* mRNA decay was determined upon treatment with 1 μg/ml of Actinomycin D (Sigma-Aldrich) for indicated time points.

### Statistical analysis

Data is presented as mean ± standard deviation. Statistical analysis was performed using GraphPad Prism 10. Statistical analysis was performed using the mean and standard deviation values. An unpaired two-sided student *t*-test was used when comparing two groups, while an ordinary one-way ANOVA test with a Tukey correction was used when comparing more than two groups. P values < 0.05 were considered significant.

## REFERENCES

1. Belz, G. T., and A. Kallies. 2010. Effector and memory CD8+ T cell differentiation: Toward a molecular understanding of fate determination. Current Opinion in Immunology 22: 279–285.

2. D’Cruz, L. M., M. P. Rubinstein, and A. W. Goldrath. 2009. Surviving the crash: Transitioning from effector to memory CD8+ T cell. Seminars in Immunology 21: 92–98.

3. Chapman, N. M., M. R. Boothby, and H. Chi. 2020. Metabolic coordination of T cell quiescence and activation. Nat Rev Immunol 20: 55–70.

4. Henning, A. N., R. Roychoudhuri, and N. P. Restifo. 2018. Epigenetic control of CD8+ T cell differentiation. Nat Rev Immunol 18: 340–356.

5. Kaech, S. M., and W. Cui. 2012. Transcriptional control of effector and memory CD8+ T cell differentiation. Nature Reviews Immunology 12: 749–761.

6. Daniels, M. A., and E. Teixeiro. 2015. TCR Signaling in T Cell Memory. Front. Immunol. 6.

7. Hwang, J.-R., Y. Byeon, D. Kim, and S.-G. Park. 2020. Recent insights of T cell receptor-mediated signaling pathways for T cell activation and development. Exp Mol Med 52: 750– 761.

8. Jurgens, A. P., B. Popović, and M. C. Wolkers. 2021. T cells at work: How post-transcriptional mechanisms control T cell homeostasis and activation. European Journal of Immunology 51: 2178–2187.

9. Salerno, F., and M. C. Wolkers. 2015. T-cells require post-transcriptional regulation for accurate immune responses. Biochemical Society Transactions 43: 1201–1207.

10. Yoshinaga, M., and O. Takeuchi. 2019. Post-transcriptional control of immune responses and its potential application. Clin Transl Immunol 8.

11. Chemin, K., C. Gerstner, and V. Malmström. 2019. Effector functions of CD4+ T cells at the site of local autoimmune inflammation-lessons from rheumatoid arthritis. Frontiers in Immunology 10: 1–15.

12. Das, D., S. Akhtar, S. Kurra, S. Gupta, and A. Sharma. 2019. Emerging role of immune cell network in autoimmune skin disorders: An update on pemphigus, vitiligo and psoriasis. Cytokine and Growth Factor Reviews 45: 35–44.

13. Killestein, J., M. J. Eikelenboom, T. Izeboud, N. F. Kalkers, H. J. Adèr, F. Barkhof, R. A. W. Van Lier, B. M. J. Uitdehaag, and C. H. Polman. 2003. Cytokine producing CD8+ T cells are correlated to MRI features of tissue destruction in MS. Journal of Neuroimmunology 142: 141–148.

14. Fara, A., Z. Mitrev, R. A. Rosalia, and B. M. Assas. 2020. Cytokine storm and COVID-19: a chronicle of pro-inflammatory cytokines. Open Biol. 10: 200160.

15. Almeida, J. R., D. A. Price, L. Papagno, Z. A. Arkoub, D. Sauce, E. Bornstein, T. E. Asher, A. Samri, A. Schnuriger, I. Theodorou, D. Costagliola, C. Rouzioux, H. Agut, A.-G. Marcelin, D. Douek, B. Autran, and V. Appay. 2007. Superior control of HIV-1 replication by CD8+ T cells is reflected by their avidity, polyfunctionality, and clonal turnover. Journal of Experimental Medicine 204: 2473–2485.

16. Ciuffreda, D., D. Comte, M. Cavassini, E. Giostra, L. Bühler, M. Perruchoud, M. H. Heim, M. Battegay, D. Genné, B. Mulhaupt, R. Malinverni, C. Oneta, E. Bernasconi, M. Monnat, A. Cerny, C. Chuard, J. Borovicka, G. Mentha, M. Pascual, J.-J. Gonvers, G. Pantaleo, and V. Dutoit. 2008. Polyfunctional HCV-specific T-cell responses are associated with effective control of HCV replication. Eur. J. Immunol. 38: 2665–2677.

17. Zhang, B., T. Karrison, D. A. Rowley, and H. Schreiber. 2008. IFN-γ– and TNF-dependent bystander eradication of antigen-loss variants in established mouse cancers. J. Clin. Invest. 118: 1398–1404.

18. Patel, S. J., N. E. Sanjana, R. J. Kishton, A. Eidizadeh, S. K. Vodnala, M. Cam, J. J. Gartner, L. Jia, S. M. Steinberg, T. N. Yamamoto, A. S. Merchant, G. U. Mehta, A. Chichura, O. Shalem, E. Tran, R. Eil, M. Sukumar, E. P. Guijarro, C.-P. Day, P. Robbins, S. Feldman, G. Merlino, F. Zhang, and N. P. Restifo. 2017. Identification of essential genes for cancer immunotherapy. Nature 548: 537–542.

19. Nicolet, B. P., N. D. Zandhuis, V. M. Lattanzio, and M. C. Wolkers. 2021. Sequence determinants as key regulators in gene expression of T cells. Immunological reviews 1–20.

20. Gagnon, J. D., and K. M. Ansel. 2019. MicroRNA regulation of CD8+ T cell responses. Non-coding RNA Investigation 3: 24–24.

21. Salerno, F., M. Turner, and M. C. Wolkers. 2020. Dynamic Post-Transcriptional Events Governing CD8+ T Cell Homeostasis and Effector Function. Trends in Immunology 41: 240– 254.

22. Gerstberger, S., M. Hafner, and T. Tuschl. 2014. A census of human RNA-binding proteins. Nature Reviews Genetics 15: 829–845.

23. Petkau, G., T. J. Mitchell, K. Chakraborty, S. E. Bell, V. D’Angeli, L. Matheson, D. J. Turner, A. Saveliev, O. Gizlenci, F. Salerno, P. D. Katsikis, and M. Turner. 2022. The timing of differentiation and potency of CD8 effector function is set by RNA binding proteins. Nature Communications 13.

24. Galloway, A., H. Ahlfors, M. Turner, L. S. Bell, and K. U. Vogel. 2016. The RNA-Binding Proteins Zfp36l1 and Zfp36l2 Enforce the Thymic β-Selection Checkpoint by Limiting DNA Damage Response Signaling and Cell Cycle Progression. The Journal of Immunology 197: 2673–2685.

25. Moore, M. J., N. E. Blachere, J. J. Fak, C. Y. Park, K. Sawicka, S. Parveen, I. Zucker-Scharff, B. Moltedo, A. Y. Rudensky, and R. B. Darnell. 2018. ZFP36 RNA-binding proteins restrain T cell activation and anti-viral immunity. eLife 7: 920–926.

26. Cook, M. E., T. R. Bradstreet, A. M. Webber, J. Kim, A. Santeford, K. M. Harris, M. K. Murphy, J. Tran, N. M. Abdalla, E. A. Schwarzkopf, S. C. Greco, C. M. Halabi, R. S. Apte, P. J. Blackshear, and B. T. Edelson. 2022. The ZFP36 family of RNA binding proteins regulates homeostatic and autoreactive T cell responses. 0981.

27. Petkau, G., T. J. Mitchell, M. J. Evans, L. Matheson, F. Salerno, and M. Turner. 2023. Zfp36l1 establishes the high affinity CD8 T[cell response by directly linking TCR affinity to cytokine sensing. Eur J Immunol 2350700.

28. Hudson, B. P., M. A. Martinez-Yamout, H. J. Dyson, and P. E. Wright. 2004. Recognition of the mRNA AU-rich element by the zinc finger domain of TIS11d. Nat Struct Mol Biol 11: 257–264.

29. Blackshear, P. J., W. S. Lai, E. A. Kennington, G. Brewer, G. M. Wilson, X. Guan, and P. Zhou. 2003. Characteristics of the Interaction of a Synthetic Human Tristetraprolin Tandem Zinc Finger Peptide with AU-rich Element-containing RNA Substrates. Journal of Biological Chemistry 278: 19947–19955.

30. Hodson, D. J., M. L. Janas, A. Galloway, S. E. Bell, S. Andrews, C. M. Li, R. Pannell, C. W. Siebel, H. R. MacDonald, K. De Keersmaecker, A. A. Ferrando, G. Grutz, and M. Turner. 2010. Deletion of the RNA-binding proteins ZFP36L1 and ZFP36L2 leads to perturbed thymic development and T lymphoblastic leukemia. Nature Immunology 11: 717–724.

31. Ogilvie, R. L., M. Abelson, H. H. Hau, I. Vlasova, P. J. Blackshear, and P. R. Bohjanen. 2005. Tristetraprolin Down-Regulates *IL-2* Gene Expression through AU-Rich Element-Mediated mRNA Decay. The Journal of Immunology 174: 953–961.

32. Lee, H. H., N. A. Yoon, M.-T. Vo, C. W. Kim, J. M. Woo, H. J. Cha, Y. W. Cho, B. J. Lee, W. J. Cho, and J. W. Park. 2012. Tristetraprolin down-regulates IL-17 through mRNA destabilization. FEBS Letters 586: 41–46.

33. Matheson, L. S., G. Petkau, B. Sáenz-Narciso, V. D’Angeli, J. McHugh, R. Newman, H. Munford, J. West, K. Chakraborty, J. Roberts, S. Łukasiak, M. D. Díaz-Muñoz, S. E. Bell, S. Dimeloe, and M. Turner. 2022. Multiomics analysis couples mRNA turnover and translational control of glutamine metabolism to the differentiation of the activated CD4+ T cell. Sci Rep 12: 19657.

34. Popović, B., B. P. Nicolet, A. Guislain, S. Engels, A. P. Jurgens, N. Paravinja, J. J. Freen-van Heeren, F. P. J. Van Alphen, M. Van Den Biggelaar, F. Salerno, and M. C. Wolkers. 2023. Time-dependent regulation of cytokine production by RNA binding proteins defines T cell effector function. Cell Reports 42: 112419.

35. Salerno, F., S. Engels, M. van den Biggelaar, F. P. J. van Alphen, A. Guislain, W. Zhao, D. L. Hodge, S. E. Bell, J. P. Medema, M. von Lindern, M. Turner, H. A. Young, and M. C. Wolkers. 2018. Translational repression of pre-formed cytokine-encoding mRNA prevents chronic activation of memory T cells. Nature Immunology 19: 828–837.

36. Alison Galloway, Alexander Saveliev, Sebastian Łukasiak, Daniel J. Hodson, Daniel Bolland, Kathryn Balmanno, Helena Ahlfors, 1 Elisa Monzón-Casanova, Sara Ciullini Mannurita, Lewis S. Bell, Simon Andrews, Manuel D. Díaz-Muñoz, Simon J. Cook, Anne Corcoran, M. T. 2016. RNA-binding proteins ZFP36L1 and ZFP36L2 promote cell quiescence. Science 352: 453–459.

37. Weiss, J. M., A. M. Bilate, M. Gobert, Y. Ding, M. A. Curotto De Lafaille, C. N. Parkhurst, H. Xiong, J. Dolpady, A. B. Frey, M. G. Ruocco, Y. Yang, S. Floess, J. Huehn, S. Oh, M. O. Li, R. E. Niec, A. Y. Rudensky, M. L. Dustin, D. R. Littman, and J. J. Lafaille. 2012. Neuropilin 1 is expressed on thymus-derived natural regulatory T cells, but not mucosa-generated induced Foxp3+ T reg cells. Journal of Experimental Medicine 209: 1723–1742.

38. Van Stipdonk, M. J. B., E. E. Lemmens, and S. P. Schoenberger. 2001. Naïve CTLs require a single brief period of antigenic stimulation for clonal expansion and differentiation. Nat Immunol 2: 423–429.

39. Salerno, F., A. Guislain, D. Cansever, and M. C. Wolkers. 2016. TLR-Mediated Innate Production of IFN-γ by CD8+ T Cells Is Independent of Glycolysis. The Journal of Immunology 196: 3695–3705.

40. Masopust, D., V. Vezys, A. L. Marzo, and L. Lefrançois. 2001. Preferential Localization of Effector Memory Cells in Nonlymphoid Tissue. Science 291: 2413–2417.

41. White, J. T., E. W. Cross, M. A. Burchill, T. Danhorn, M. D. McCarter, H. R. Rosen, B. O’Connor, and R. M. Kedl. 2016. Virtual memory T cells develop and mediate bystander protective immunity in an IL-15-dependent manner. Nature Communications 7: 11291.

42. Marusina, A. I., Y. Ono, A. A. Merleev, M. Shimoda, H. Ogawa, E. A. Wang, K. Kondo, L. Olney, G. Luxardi, Y. Miyamura, T. D. Yilma, I. B. Villalobos, J. W. Bergstrom, D. G. Kronenberg, A. M. Soulika, I. E. Adamopoulos, and E. Maverakis. 2017. CD4+ virtual memory: Antigen-inexperienced T cells reside in the naïve, regulatory, and memory T cell compartments at similar frequencies, implications for autoimmunity. Journal of Autoimmunity 77: 76–88.

43. Salerno, F., N. A. Paolini, R. Stark, M. von Lindern, and M. C. Wolkers. 2017. Distinct PKC-mediated posttranscriptional events set cytokine production kinetics in CD8 ^+^ T cells. Proceedings of the National Academy of Sciences 114: 201704227.

44. De Witte, M. A., M. Coccoris, M. C. Wolkers, M. D. Van Den Boom, E. M. Mesman, J.-Y. Song, M. Van Der Valk, J. B. A. G. Haanen, and T. N. M. Schumacher. 2006. Targeting self-antigens through allogeneic TCR gene transfer. Blood 108: 870–877.

45. Thommen, D. S., V. H. Koelzer, P. Herzig, A. Roller, M. Trefny, S. Dimeloe, A. Kiialainen, J. Hanhart, C. Schill, C. Hess, S. S. Prince, M. Wiese, D. Lardinois, P. C. Ho, C. Klein, V. Karanikas, K. D. Mertz, T. N. Schumacher, and A. Zippelius. 2018. A transcriptionally and functionally distinct pd-1 + cd8 + t cell pool with predictive potential in non-small-cell lung cancer treated with pd-1 blockade. Nature Medicine 24.

46. Ahmadzadeh, M., L. A. Johnson, B. Heemskerk, J. R. Wunderlich, M. E. Dudley, D. E. White, and S. A. Rosenberg. 2009. Tumor antigen–specific CD8 T cells infiltrating the tumor express high levels of PD-1 and are functionally impaired. Blood 114: 1537–1544.

47. Salerno, F., A. Guislain, J. J. Freen-Van Heeren, B. P. Nicolet, H. A. Young, and M. C. Wolkers. 2019. Critical role of post-transcriptional regulation for IFN-γ in tumor-infiltrating T cells. OncoImmunology 8: 1–12.

48. Xu, Y., W. Wu, Q. Han, Y. Wang, C. Li, P. Zhang, and H. Xu. 2019. Post-translational modification control of RNA-binding protein hnRNPK function. Open Biol. 9: 180239.

49. Sternburg, E. L., L. A. Gruijs Da Silva, and D. Dormann. 2022. Post-translational modifications on RNA-binding proteins: accelerators, brakes, or passengers in neurodegeneration? Trends in Biochemical Sciences 47: 6–22.

50. Clark, A. R., and J. L. E. Dean. 2016. The control of inflammation via the phosphorylation and dephosphorylation of tristetraprolin: A tale of two phosphatases. Biochemical Society Transactions 44: 1321–1337.

51. Navarro, M. N., J. Goebel, C. Feijoo-Carnero, N. Morrice, and D. A. Cantrell. 2011. Phosphoproteomic analysis reveals an intrinsic pathway for the regulation of histone deacetylase 7 that controls the function of cytotoxic T lymphocytes. Nature Immunology 12: 352–362.

52. Tan, H., K. Yang, Y. Li, T. I. Shaw, Y. Wang, D. B. Blanco, X. Wang, J.-H. Cho, H. Wang, S. Rankin, C. Guy, J. Peng, and H. Chi. 2017. Integrative Proteomics and Phosphoproteomics Profiling Reveals Dynamic Signaling Networks and Bioenergetics Pathways Underlying T Cell Activation. Immunity 46: 488–503.

53. Brenes, A. J., A. I. Lamond, and D. A. Cantrell. 2023. The Immunological Proteome Resource. Nat Immunol 24: 731–731.

54. Rouvière, J. O., M. Bulfoni, A. Tuck, B. Cosson, F. Devaux, and B. Palancade. 2018. A SUMO-dependent feedback loop senses and controls the biogenesis of nuclear pore subunits. Nat Commun 9: 1665.

55. Campagnaro, G. D., E. Nay, M. J. Plevin, A. K. Cruz, and P. B. Walrad. 2021. Arginine Methyltransferases as Regulators of RNA-Binding Protein Activities in Pathogenic Kinetoplastids. Front. Mol. Biosci. 8: 692668.

56. Fabian, M. R., F. Frank, C. Rouya, N. Siddiqui, W. S. Lai, A. Karetnikov, P. J. Blackshear, B. Nagar, and N. Sonenberg. 2013. Structural basis for the recruitment of the human CCR4–NOT deadenylase complex by tristetraprolin. Nat Struct Mol Biol 20: 735–739.

57. Zandhuis, N. D., B. P. Nicolet, and M. C. Wolkers. 2021. RNA-Binding Protein Expression Alters Upon Differentiation of Human B Cells and T Cells. Frontiers in Immunology 12.

58. Dighe, A. S., E. Richards, L. J. Old, and R. D. Schreiber. 1994. Enhanced in vivo growth and resistance to rejection of tumor cells expressing dominant negative IFNγ receptors. Immunity 1: 447–456.

59. Nastala, C. L., H. D. Edington, T. G. McKinney, H. Tahara, M. A. Nalesnik, M. J. Brunda, M. K. Gately, S. F. Wolf, R. D. Schreiber, and W. J. Storkus. 1994. Recombinant IL-12 administration induces tumor regression in association with IFN-gamma production. The Journal of Immunology 153: 1697–1706.

60. Kemna, J., E. Gout, L. Daniau, J. Lao, K. Weißert, S. Ammann, R. Kühn, M. Richter, C. Molenda, A. Sporbert, D. Zocholl, R. Klopfleisch, H. Lortat-Jacob, P. Aichele, T. Kammertoens, and T. Blankenstein. 2023. IFNγ binding to extracellular matrix prevents fatal systemic toxicity. Nat Immunol .

61. Vredevoogd, D. W., T. Kuilman, M. A. Ligtenberg, J. Boshuizen, K. E. Stecker, B. de Bruijn, O. Krijgsman, X. Huang, J. C. N. Kenski, R. Lacroix, R. Mezzadra, R. Gomez-Eerland, M. Yildiz, I. Dagidir, G. Apriamashvili, N. Zandhuis, V. van der Noort, N. L. Visser, C. U. Blank, M. Altelaar, T. N. Schumacher, and D. S. Peeper. 2019. Augmenting Immunotherapy Impact by Lowering Tumor TNF Cytotoxicity Threshold. Cell 1–15.

62. Aktas, E., U. C. Kucuksezer, S. Bilgic, G. Erten, and G. Deniz. 2009. Relationship between CD107a expression and cytotoxic activity. Cellular Immunology 254: 149–154.

63. Hurkmans, D. P., E. A. Basak, N. Schepers, E. Oomen-De Hoop, C. H. Van Der Leest, S. El Bouazzaoui, S. Bins, S. L. W. Koolen, S. Sleijfer, A. A. M. Van Der Veldt, R. Debets, R. H. N. Van Schaik, J. G. J. V. Aerts, and R. H. J. Mathijssen. 2020. Granzyme B is correlated with clinical outcome after PD-1 blockade in patients with stage IV non-small-cell lung cancer. J Immunother Cancer 8: e000586.

64. Gocher, A. M., C. J. Workman, and D. A. A. Vignali. 2022. Interferon-γ: teammate or opponent in the tumour microenvironment? Nat Rev Immunol 22: 158–172.

65. Hogquist, K. A., S. C. Jameson, W. R. Heath, J. L. Howard, M. J. Bevan, and F. R. Carbone. 1994. T cell receptor antagonist peptides induce positive selection. Cell 76: 17–27.

66. Morita, S., T. Kojima, and T. Kitamura. 2000. Plat-E: an efficient and stable system for transient packaging of retroviruses. Gene Ther 7: 1063–1066.

67. Refaeli, Y., L. Van Parijs, S. I. Alexander, and A. K. Abbas. 2002. Interferon γ Is Required for Activation-induced Death of T Lymphocytes. Journal of Experimental Medicine 196: 999–1005.

68. Heinen, A. P., F. Wanke, S. Moos, S. Attig, H. Luche, P. P. Pal, N. Budisa, H. J. Fehling, A. Waisman, and F. C. Kurschus. 2014. Improved method to retain cytosolic reporter protein fluorescence while staining for nuclear proteins. Cytometry Part A 85: 621–627.

69. Salerno, F., A. Guislain, D. Cansever, and M. C. Wolkers. 2016. TLR-Mediated Innate Production of IFN-γ by CD8 + T Cells Is Independent of Glycolysis. The Journal of Immunology 196: 3695–3705.

